# Unraveling the interaction between doxorubicin and DNA origami nanostructures for customizable chemotherapeutic drug release

**DOI:** 10.1101/2020.05.13.088054

**Authors:** Heini Ijäs, Boxuan Shen, Amelie Heuer-Jungemann, Adrian Keller, Mauri A. Kostiainen, Tim Liedl, Janne A. Ihalainen, Veikko Linko

## Abstract

Doxorubicin (DOX) is a common drug in cancer chemotherapy, and its high DNA-binding affinity can be harnessed in preparing DOX-loaded DNA nanostructures for targeted delivery and therapeutics. Although DOX has been widely studied, the existing literature of DOX-loaded DNA-carriers remains limited and incoherent. Here, based on an in-depth spectroscopic analysis, we characterize and optimize the DOX loading into different 2D and 3D scaffolded DNA origami nanostructures (DONs). In our experimental conditions, all DONs show similar DOX binding capacities (one DOX molecule per two to three base pairs), and the binding equilibrium is reached within seconds, remarkably faster than previously acknowledged. To characterize drug release profiles, DON degradation and DOX release from the complexes upon DNase I digestion was studied. For the employed DONs, the relative doses (DOX molecules released per unit time) may vary by two orders of magnitude depending on the DON superstructure. In addition, we identify DOX aggregation mechanisms and spectral changes linked to pH, magnesium, and DOX concentration. These features have been largely ignored in experimenting with DNA nanostructures, but are probably the major sources of the incoherence of the experimental results so far. Therefore, we believe this work can act as a guide to tailoring the release profiles and developing better drug delivery systems based on DNA-carriers.

## Introduction

The possibility to employ DNA molecules in engineering artificial nanostructures (1, 2) has drawn increasing attention during the past two decades (3–5). The intense development of DNA nanotechnology has yielded new methods to build user-defined nano-objects (6), such as DNA origami (7–11), for a variety of scientific and technological uses (12–16). In particular, these custom DNA nanoshapes show considerable promise in biomedicine and drug delivery (17–21). Rationally designed DNA nanovehicles can encapsulate and display selected cargoes (22–25), act as therapeutics themselves (26), serve as platforms for various targeting ligands and tailored nucleic acid sequences (27, 28), or directly host diverse DNA-binding drugs (29, 30). In the latter case, the most frequently used drug is anthracycline doxorubicin (DOX), a fluorescent DNA-intercalator, which is applied in the treatments of several cancer types and primarily in solid tumor growth suppression (31). Its main mechanism of action takes place *via* type IIA DNA topoisomerase inhibition, but it also affects multiple other cellular processes through DNA intercalation and generation of reactive oxygen species (ROS) (32). The therapeutic potency of various DOX-loaded DNA origami nanostructures (DONs) has been demonstrated using *in vitro* and in vivo models in a number of reports (33–43).

The presumed intercalation and release of DOX are typically characterized using spectroscopic indicators such as spectral changes of visible light absorption or DOX fluorescence quenching upon DNA binding. However, besides intercalation, DOX may be complexed with DNA through (pre-intercalation) minor-groove binding and stacking into aggregates depending on the DNA sequence, prevalent DOX concentration and experimental conditions such as pH or the ionic strength of the solution (44–46). Spectroscopic features of bound DOX are likewise dependent on the mode of interaction. In addition, DOX molecules have two distinct protonation states within a physiologically relevant pH range (pH ~4–9) and they are prone to self-association at high concentrations (47). Therefore, spectroscopic properties of DOX are also subject to change in different media compositions. These effects need to be carefully differentiated from the changes induced by DNA binding to avoid misleading interpretations of DOX loading capacity, release efficiency and the therapeutic effect (20).

In this work, we systematically study the binding of DOX to five structurally distinct two- (2D) and three-dimensional (3D) DONs (one exemplary structure shown in Figure 1). By means of absorption and fluorescence spectroscopy, we optimize the loading process and uncover the contributions of the ionic strength, pH, and DOX concentration. The obtained results reveal that the DOX binding capacity of DONs has often been substantially overestimated, in some previously reported cases by more than two orders of magnitude.

**Figure 1.**
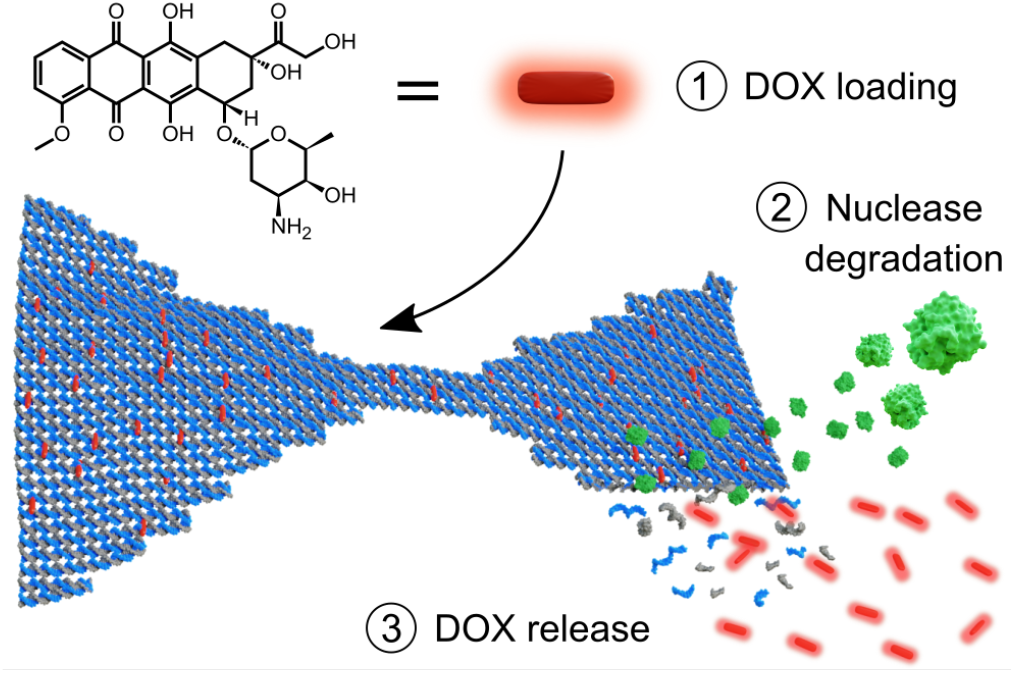
Schematic of the doxorubicin (DOX) loading into a DNA origami nanostructure (DON) and subsequent release upon enzymatic degradation. Here, we 1) study how DOX is loaded into DONs (in seconds), optimize the conditions for the loading by monitoring the spectroscopic features of DOX, and characterize the formed DOX-DON complexes. Through simultaneous real-time detection of the absorbance and fluorescence changes of the DOX-loaded DONs, we then 2) monitor the degradation of DONs into single-stranded DNA fragments by nucleases (DNase I, green) (in minutes to hours under a DNase I concentration of 34 U mL^-1^), and 3) characterize the subsequent DOX release profiles of different DONs and show that the release profiles of DOX depend on the DNA origami superstructure and the applied DOX content.

Finally, we mimic one plausible and physiologically relevant DOX release pathway by subjecting the DOX-loaded DONs to deoxyribonuclease I (DNase I) digestion (see Figure 1) (48–50). Real-time monitoring of the spectroscopic changes during the digestion show that both the DNA degradation rates and the DOX release profiles depend on the DNA origami superstructure and the amount of loaded DOX. We believe that through identification of these fundamental and some previously undiscovered features of the loading process, the spectroscopic properties of DOX, as well as the superstructure-dependent stability factors of DONs in physiological conditions (51–54), it may become possible to rationally design the delivery capability, control the dose, and thus achieve the optimal therapeutic efficacy in DOX delivery.

## Materials and methods

### Materials

Doxorubicin hydrochloride (HPLC-purified, Sigma-Aldrich) was dissolved in Milli-Q water for a 10 mM stock solution, divided into aliquots and stored at –20°C. After thawing, the stock solution was stored at +4°C and used within 1–2 days.

DNase I was purchased from Sigma-Aldrich. A stock solution was prepared at 2 U/μL concentration in deionized water and stored in aliquots at –20 °C, and after thawing stored at room temperature (RT) and used within the same day.

The staple oligonucleotides for DON folding were purchased from Integrated DNA Technologies. For the Rothe-mund triangle (7), bowtie (55), double-L (55), and the closed capsule (25), the staple strand sequences and folding protocols were adopted from the original publications. The 24-helix bundle (24HB) was designed using caDNAno (56) and its shape was predicted using CanDo software (57, 58). The design of the 24HB structure is presented in the Supplementary Figures 16–18, and its staple strand sequences in the Supplementary Table 5. The self-assembly reaction for the 24HB was carried out in a buffer containing 1× TAE and 17.5 mM MgCl_2_. The reactions were heated to 65 °C, and assembled by first cooling to 59 °C with a rate of 1 °C/15 min and then to 12 °C with rate 0.25 °C/45 min. The 7,249 nt long M13mp18 scaffold was used for folding the triangle, bowtie and double-L. The extended 7,560 nt and 8,064 nt variants were used for folding the 24HB and capsule, respectively. All DNA scaffold strands were purchased from Tilibit Nanosystems at 100 nM concentration. After thermal annealing, the DONs were purified of excess staple strands using polyethylene glycol (PEG) precipitation (59) in the presence of 7.5% (w/v) PEG 8000 (Sigma-Aldrich). After purification, the DONs were resuspended in 40 mM Tris, 10 mM MgCl_2_, pH 7.4 and incubated overnight at RT before use. The structural integrity was verified by AFM and TEM.

### Spectroscopy techniques

Unless otherwise indicated, all UV-Vis absorption and fluorescence measurements were carried out with the Aqualog absorbance-fluorescence system (Horiba Scientific) operated with the Aqualog Software (v4.2) (Horiba Scientific), with the sample in a 10 mm optical path length cuvette. In spectral scans, a 3D excitation-emission matrix and the absorption spectrum of the sample were recorded simultaneously using an excitation light scan at 2 nm increments between 240– 700 nm with 5 nm slit width. The emission spectrum for each excitation wavelength was collected between 245.16– 827.17 nm at 1.16 nm increments with the CCD array. All measurements were performed at RT.

A 3 μM DOX concentration was selected for most experiments for avoiding possible DOX aggregation and self-quenching at high concentration, but also for performing accurate spectroscopic analysis in the low absorbance (*A* < 0.1) region where both *A* and emission intensity *(I)* values exhibit linear dependency on the concentration of the studied molecules.

### Free DOX characterization: spectroscopic analysis of the effect of pH and MgCl_2_ concentration

For studying the effect of buffer pH on the spectroscopic properties of DOX, 40 mM Tris-HCl buffers at pH 6.0, 7.0, 7.4, 7.8, 8.0, 8.2, 8.6, or 9.0 were prepared by dissolving the required amount of Tris base in water and adjusting the pH of the solution with 1 M HCl. 3 μM DOX solutions were prepared in each of the buffers from the 10 mM stock solution and the UV-Vis absorption and 3D excitation-emission matrices of all samples were measured separately.

For the measurement of DOX absorption and fluorescence in the presence of different MgCl_2_ concentrations, 3 μM solutions of DOX were prepared in 40 mM Tris-HCl buffers at pH 7.4 at both 0 mM and 100 mM MgCl_2_ concentration. The absorption and fluorescence spectra of both samples were first recorded separately as described. The 3 μM DOX in the 100 mM MgCl_2_ buffer was then added in small volumes into the 0 mM MgCl_2_ DOX solution in the cuvette. After each addition, the sample was mixed by vortexing and its absorption and fluorescence spectra were recorded. The MgCl_2_ concentration at each titration step was calculated according to the added volume of the 100 mM MgCl_2_ DOX solution.

### DOX loading and self-aggregation study

The DOX-DON loading was studied in three different buffers: 1) 40 mM Tris with 10 mM MgCl_2_, pH 7.4, 2) 40 mM Tris with 10 mM MgCl_2_, pH 8.0, and 3) 1× TAE with 12.5 mM MgCl_2_ (1 × FOB), pH 8.0. DOX-DON samples were prepared in each of the studied buffers by mixing the triangle DON (at 2.5 nm final concentration) with either 2 mM, 200 μM, or 20 μM DOX. Corresponding reference samples of DOX without DONs were prepared at 2 mM, 200 μM, or 20 μM DOX concentration in each of the buffers.

The UV-Vis absorption spectra of the solutions were measured in the beginning of the experiment using either a Cytation 3 cell imaging multi-mode reader (BioTek) on a Take3 plate with a 0.5 mm optical path length (2 mM and 200 μM samples), or Nanodrop ND-1000 with a 1 mm path length (20 μM samples). After 24 h, 48 h, or 96 h incubation in dark at RT, the samples were centrifuged for 10 min at 14,000 *g* to separate the fraction of insoluble DOX. The effect of the applied centrifugal force (2,000–14,000 *g)* was additionally tested with 200 μM DOX in FOB pH 8.0 after 24 h incubation (Supplementary Methods and Supplementary Figure 3). The concentration of DOX in the supernatant was determined by removing a small volume of the supernatant and measuring the UV-Vis absorption similarly as in the beginning of the incubation. The DOX concentration in the supernatant (*c*_free_) relative to the concentration at *t* = 0 was calculated from the *A*_480_, and *c*_aggregate_ was defined as *c*_aggregate_ = *c*_0_ – *c*_free_. The experiment was repeated three times and the final values were reported as the mean ± standard error.

### DNA origami – DOX titrations

#### UV-Vis and fluorescence spectroscopy

Association of DOX with DONs was studied by titrating a solution of 3 μM DOX in 40 mM Tris, 10 mM MgCl_2_, pH 7.4 with a solution containing *ca*. 40 nM DONs (triangle, bowtie, double-L, capsule, or 24HB) and 3 μM DOX in the same buffer. 40 nM DON concentration corresponds to 558–605 μM base pair concentration [*c*(bp)] depending on the DON design (see details in Supplementary Table 1). After each addition of the titrant, the sample was mixed by vortexing and let to equilibrate for 2.5 min before measuring the absorption and fluorescence spectra.

The effect of the equilibrium time was further studied with kinetic measurements for both 2D and 3D structures. The absorption and fluoresence spectra of 3 μM DOX in 40 mM Tris, 10 mM MgCl_2_, pH 7.4 were first recorded in the absence of DONs. Triangle DONs or 24HB DONs were then added at a molar ratio of bp/DOX ≈ 2 and the absorption and fluorescence spectra of the samples were collected after 0.5, 2.5, 5, 10, 15, and 20 min of incubation.

#### Data analysis and fitting

The concentration of DNA [total nucleotide concentration *c*(nt)] at each point of the titration was determined from the DNA absorption at 260 nm (*A*_260_). As both DNA and DOX absorb light at 260 nm, the contribution of DOX absorption was removed from the obtained *A*_260_ values by subtracting the *A*_260_ of 3 μM DOX solution. *c*(nt) was then determined according to the Beer-Lambert law. The molar extinction coefficient per nucleotide (*ϵ*_260_/nt) was calculated separately for each DON with a formula adapted from Hung *et al.* (60) according to the number of unpaired nucleotides (*N*_ss_) and number of hybridized nucleotides (*N*_ds_) in the design,

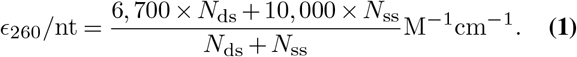

The values of *N*_ss_ and *N*_ds_ for each DON are presented in Supplementary Table 1. The value of *N*_ds_ takes into account both the base pairs formed by hybridization of the scaffold and staple strands, and the base pairs formed as secondary structures through hybridization of self-complementary regions of unpaired scaffold loops. The number of base pairs in the secondary structures was simulated with the NUPACK web application (61).

As the intercalation of DOX into DONs depends on the *ND*_s_, only the concentration of hybridized nucleotides (base pair concentration *c*(bp)_0_ = 0.5 × *c*(nt) × *N*_ds_/(*N*_ds_ + *N*_ss_) was taken into account in the analysis. To justify this, DOX was also titrated with ssDNA (Supplementary Figure 9). ss-DNA quenches DOX fluorescence only slightly when compared to the quenching caused by double-stranded DNA (dsDNA). Although all DONs used in this work, except the triangle, contain ssDNA regions at both poly-T_8_ extensions at the helix ends and at unpaired scaffold loops, their contribution to DOX quenching is thus negligible. In addition, the fraction of unpaired nucleotides (mostly the inert poly-T sequences) of the total number of nucleotides [*N*_SS_/(*N*_ds_ + *N*_ss_)] in the structures is small (0.4–12%).

For fitting the fluorescence data, emission intensity values from 450, 460, 470, 480, and 494 nm excitation were obtained from the integrated emission spectra. The values were corrected for the extinction coefficient decrease during titration by dividing with (1 — *T*) of the excitation wavelength (*T* denotes the transmittance, obtained from the simultaneous absorption measurement). The corrected values thus represent the decrease of DOX fluorescence quantum yield (Φ) upon DNA binding. The obtained values for ΔΦ_obs_ = Φ_obs_ — Φ_0_ for each *c*(bp)_0_ were fitted with a 1:2 host-guest binding model using the BindFit online fitting tool at https://supramolecular.org (62). The fit describes the change of the studied physical property (Δ*F*_obs_) as

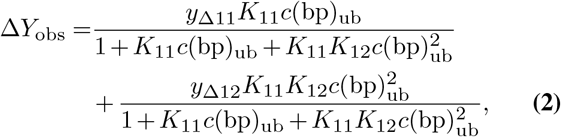

where in the case of Φ, Δ*Y*_obs_ refers to the measured ΔΦ_obs_, while *y*_Δ11_ and *y*_Δ12_ refer to the differences of the quantum yields of the 1:1 and 1:2 DOX:base pair complexes and the quantum yield of free DOX (Φ_Δ11_ and Φ_Δ12_), respectively. The binding constants *K*_11_ and *K*_12_, as well as Φ_Δ11_ and Φ_Δ12_ were obtained from the fit. In Equation 2, *c*(bp)_ub_ is the concentration of unbound base pairs, *i.e.* the free base pairs not bound to DOX, obtained from

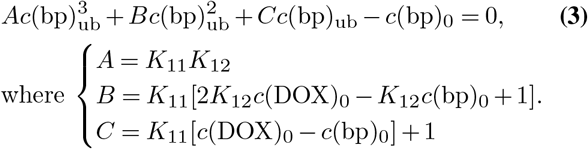

The fraction of bound DOX molecules *f*_b_ at each step of the titration was then calculated as

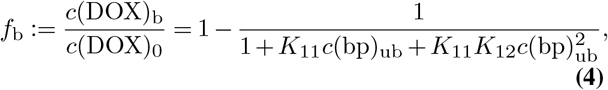

where *c*(DOX)_b_ is the calculated bound DOX concentration at the specific *c*(bp)_ub_, and *c*(DOX)_0_ denotes the total DOX concentration (3 μM).

After *K*_11_, *K*_12_, and *c*(bp)_ub_ for each *c*(bp)_0_ were obtained from the analysis of the fluorescence data, the molar extinction coefficients of the two DOX-DNA complexes, *ϵ*_11_, and *ϵ*_12_ at wavelengths 450,460,470,480, and 494 nm, were determined with non-linear least-squares curve fitting with MATLAB R2015b to the Equation 2. For absorbance, *y*_Δ11_ refers to *ϵ*_Δ11_, and *y_Δ12_* refers to *ϵ*_Δ12_ – the differences between the molar extinction coefficients of the two complexes and the molar extinction coefficient of free DOX (listed in the Supplementary Table 2). Δ*A*_obs_ is the difference between the absorbance of the sample and the absorbance of 3 μM DOX in the absence of DNA.

### DNase I digestion of DNA origami

#### Kinetic UV-Vis and fluorescence measurements

The DNase I digestion and DOX release rates of the studied DONs were determined based on the absorbance and fluorescence spectra of the samples collected during DNase I digestion. The digestion of each DON was studied both in the absence of DOX and after loading the DONs with DOX. The samples without DOX contained 2 nM DONs in 40 mM Tris, 10 mM MgCl_2_, pH 7.4. The DON-DOX samples contained 2 nM DONs and either 3 or 6 μM DOX. Unbound DOX in the solution was not removed before the digestion, as the removal of free DOX from the system would disturb the binding equilibrium and promote the dissociation of bound DOX from the DONs, complicating the analysis of the release caused solely by DNase I digestion. In addition, the comparison between samples can be performed more accurately when the concentration of each component in the sample (DNA, bound DOX, and free DOX) is known precisely and not changed with purification protocols, such as spin-filtration.

The absorbance and fluorescence spectra of the samples were first collected without DNase I. DNase I was then added to final concentration of 34 U mL^-1^, the sample was gently mixed with a pipette, and the absorbance and fluorescence spectra were collected at regular time intervals until the digestion had been completed. The total duration of the experiment was adjusted for each DON, ranging from < 1 h required for a full digestion of the triangle DON with 0 μM DOX, to > 40 h for the 24HB with 6 μM DOX.

Samples containing only 3 μM or 6 μM DOX and DNase I were measured similarly for obtaining references for the fluorescence quantum yield of free DOX in the same experimental conditions.

#### Data analysis of the kinetic measurements

The nuclease digestion of the DONs was quantified from the *A_260_* value, which increases during the digestion. The percentage of intact dsDNA residues (% intact) at each time point t was calculated as

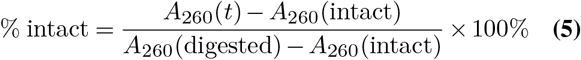

where *A*_260_(*t*) is the *A*_260_ value detected at a time point *t, A*_260_ (intact) is the *A*_260_ of fully intact structures measured before addition of DNase I, and *A*_260_ (digested) is the *A*_260_ of fully digested structures, measured after the digestion has been completed (*A*_260_ value stabilized). The DNase I digestion rates were determined by fitting a linear regression to the obtained % intact values vs. time with MATLAB R2015b at the initial period of the digestion where the *A*_260_ increases linearly. Digestion rates were determined from three repeated experiments and reported as the mean ± standard error.

The analysis of DOX release from the DONs during DNase I digestion was based on the recovery of DOX quantum yield (fluorescence emission intensity from 494 nm excitation divided by (1 – *T*)). The mole fraction of bound DOX molecules at each time point [*f*_b_ (*t*)] was calculated with Equation 4. In Equation 4, *f*b(*t*) calculation is based on the *c*(bp)_ub_ at each time point; *c*(bp)_ub_ was solved from a modified and rearranged version of Equation 2,

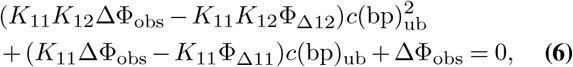

where ΔΦ_obs_ is the measured difference of the quantum yield of the DOX-DON sample and the free DOX reference of the same DOX concentration. Φ_Δ11_ and Φ_Δ12_ are the differences between the quantum yield of free DOX and the quantum yields of the 1:1 and 1:2 DOX-DNA complexes, respectively. *K*_11_, *K*_12_, Φ_Δ11_, and Φ_Δ12_ are fit parameters averaged from the titration experiments of all the studied DONs (Supplementary Table 2). The percentage of released DOX (Figure 5a) was then defined as

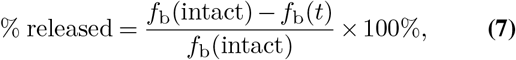

where *f*_b_(intact) is *f*_b_ of the sample before addition of DNase I. DOX release rates were acquired by fitting a linear regression to the % released vs. time in the initial linear phase of drug release. For calculating the DOX release rate in terms of relative dose (number of released molecules) per unit of time, the rate values were multiplied by the DOX concentration bound to the DONs in the intact state. The DOX release rates were determined from three repeated experiments and reported as the mean ± standard error.

### Microscopy imaging

#### Atomic force microscopy (AFM)

To prepare the 2D DON (triangle, bowtie and double-L) samples for AFM imaging, 10 μL of 3 nM DON solution in 1 × FOB (with 10 mM MgCl_2_) was pipetted onto a fresh-cleaved mica sheet and incubated for 5 min. Then the mica substrate was washed 3 times with 100 μL ddH_2_O by allowing it to flow from one end of the mica to the other and being blotted using a piece of cleanroom sheet. Finally, the sample was rigorously dried by pressurized N2 gas. The AFM imaging was carried out using either a JPK NanoWizard ULTRA Speed with USCF0.3-k0.3 cantilevers (NanoWorld) in a liquid cell filled with 1 × FOB (with 10 mM MgCl_2_) or using a Bruker Dimension Icon instrument in Scanasyst air mode and Scanasyst-air probes.

#### Transmission electron microscopy (TEM)

The 3D DON samples (capsule and 24HB) were characterized using a Tecnai T12 TEM instrument. Copper TEM grids with both carbon and formvar films (FCF400-Cu from Electron Microscopy Sciences) were cleaned with O_2_ plasma for 20 s, followed by pipetting 3 μL of 20 nM DON solution on the grid and incubating for 2 min. Then the excess amount of solution was blotted with a piece of filter paper. To achieve better contrast, the sample was immediately stained with 20 μL of 2% uranyl formate for 40 s followed by blotting the staining solution with the filter paper. The grid was let to dry for at least 30 min before imaging.

## Results

### Effects of buffer conditions on the spectroscopic features of DOX

To ensure that the obtained spectroscopic changes in later experiments are associated reliably with the DOX-DNA binding events and not caused by the environment, we first identified the effects of the buffer conditions on the spectroscopic properties of DOX. We performed a series of measurements on DOX in the absence of DNA in Tris-based buffers typically applied in DON experiments. In particular, we screened the effect of two buffer parameters; pH and MgCl_2_ concentration.

#### Buffer pH

For identifying the effects of buffer pH on the spectroscopic features of DOX, 40 mM Tris-HCl buffers were prepared at pH 6.0–9.0 and the absorption and fluorescence spectra of DOX were collected at each pH. The shape of the DOX absorption spectrum as well as its molar extinction coefficient (*ϵ*) depends heavily on buffer pH (Figure 2a). Between pH 6.0–8.0, the shape of the spectrum is maintained, but *ϵ* increases with decreasing pH throughout the whole absorption spectrum. For instance, *ϵ*_494_ is ~65%. higher at pH 6.0 than at pH 8.0. A higher emission intensity is also observed at lower pH values, as shown in Figure 2a inset with a 494 nm excitation.

**Figure 2.**
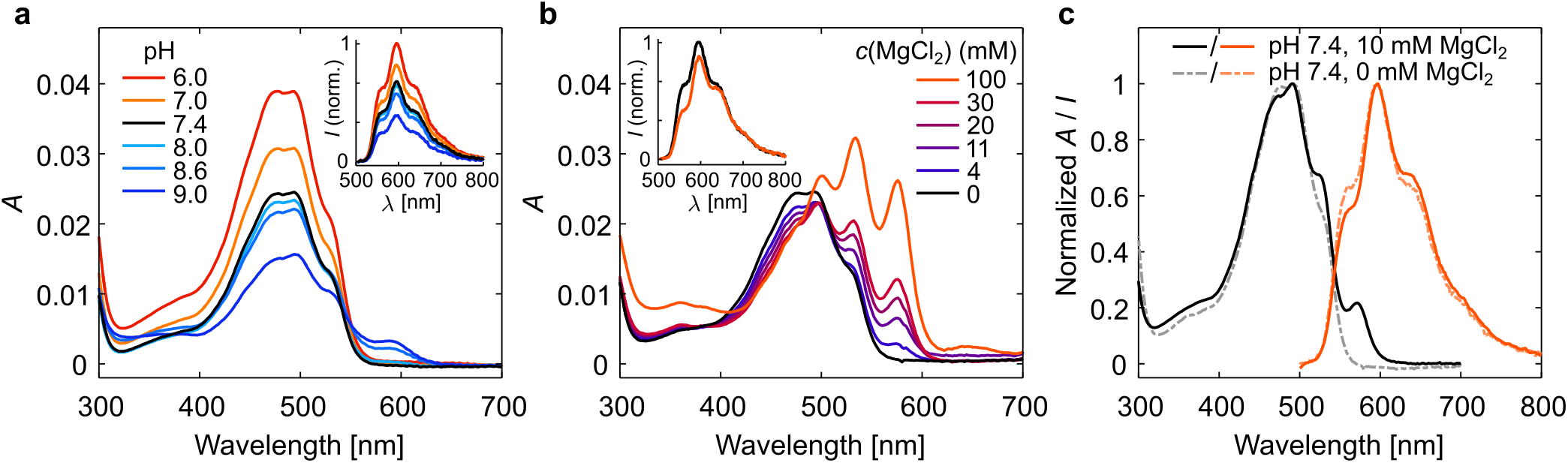
Effects of buffer conditions on the spectral features of DOX in the absence of DNA. **a** Absorption and emission (inset) spectra of 3 μM DOX in 40 mM Tris, 0 mM MgCl_2_ at pH 6.0–9.0. The emission spectra were obtained at an excitation wavelength of 494 nm. **b** Spectral features of 3 μM DOX in 40 mM Tris, pH 7.4 buffer at different MgCl_2_ concentrations. The inset figure shows a comparison of the emission spectra of the 0 mM and 100 mM samples, with the maximum emission intensity of the 0 mM MgCl_2_ sample normalized to 1. The 0 mM MgCl_2_ spectrum (black) corresponds to the pH 7.4 spectrum in Figure a. **c** Absorption (black/gray lines) and emission (orange lines) spectra of 3 μM DOX in the chosen experimental conditions – 40 mM Tris, 10 mM MgCl_2_, pH 7.4. The effect of the 10 mM MgCl_2_ concentration is shown by comparing the spectra measured at 10 mM MgCl_2_ (solid lines) with spectra measured at 0 mM MgCl_2_ (dashed lines).

Above pH 8.0, the shape of the absorption spectrum changes and a new absorption peak emerges at approximately 590 nm. Exciting the molecules at this wavelength does not lead to DOX fluorescence, thus showing that at pH 8.0 and above, an increasing fraction of DOX molecules is non-fluorescent. DOX is known to have a p*K*A value for the deprotonation of the amino sugar NH‡ group at pH 8.2 (47). The observed emergence of non-fluorescent molecules takes place around the same pH value, being thus likely associated with the deprotonation events. These observations are also in line with previous reports of DOX absorbance in high pH buffers (63), and the spectral changes could thus be expected to become even more pronounced at pH values above 9.0.

Near the p*K*_a_, the sample contains a distribution of charged and neutral molecules, and in the spectroscopic means, a mixture of fluorescent and non-fluorescent DOX molecules. While the sample is thus heterogeneous, the emission spectrum remains homogeneous as the non-fluorescent molecules do not contribute to the signal (Supplementary Figure 1). As the sample heterogeneity would nevertheless complicate the interpretation of experimental results, it is beneficial to conduct experiments at pH well below the p*K*_a_. Based on both the existing literature and the obtained spectra, an optimal pH range for further experiments was determined as 6.0–7.8, where altering the pH does not change the shape of the absorption spectrum.

#### Buffer MgCl_2_ concentration at pH 7.4

DOX is known to form complexes with metal ions, such as Fe^3+^, Cu^2+^, Mn^2+^, Ni^2+^, Co^2+^, Mg^2+^, and Zn^2+^ (63–65). Metal ion complexation thus presents another source of DOX heterogeneity in buffers supplemented with divalent cations. When the MgCl_2_ concentration in the buffer increases, both the absorption and fluorescence properties of DOX change indicating complexation of DOX with Mg^2+^ ions (Figure 2b). In the presence of 100 mM MgCl_2_, three distinct peaks at 500 nm, 534 nm, and 576 nm are observed in the absorption spectrum. The 576 nm peak emerges only in the presence of MgCl_2_, and excitation at this absorption peak leads to a fluorescence spectral shape that is rather distinct from that of DOX in the absence of MgCl_2_ (Supplementary Figure 2). While the emission spectrum of DOX at 0 mM MgCl_2_ is homogeneous over the full absorption spectrum, the addition of MgCl_2_ induces heterogeneity in the emission measurement reflected as the shape of the emission spectrum changing with the excitation wavelength (Supplementary Figure 2). As a result, the shape of the emission spectrum upon 494 nm excitation depends slightly on the MgCl_2_ concentration (Figure 2b inset).

A comparison of the absorption and fluorescence spectra of 3 μM DOX in 40 mM Tris, pH 7.4 with either 0 mM or 10 mM MgCl_2_ is shown in Figure 2c. The spectral differences indicate that at 10 mM MgCl_2_, the sample and its absorption and fluorescence spectra are a combination of pure DOX and a small concentration of the DOX-Mg^2+^ complex. Despite the slight DOX heterogeneity in these conditions, 40 mM Tris at pH 7.4 supplemented with 10 mM MgCl_2_ was chosen for all the experiments to maintain structural stability and integrity of the DONs.

### DOX loading

#### DOX self-aggregation during loading

As the buffer pH and MgCl_2_ concentration have considerable effects on the physical and spectroscopic properties of free DOX, they can be assumed to affect the association of DOX with DONs; *i.e.* the loading process. To study this, we loaded the triangle DON (Figure 4a) with DOX in three different buffers: in 40 mM Tris, 10 mM MgCl_2_ at pH 7.4 (Tris/Mg^2+^ pH 7.4); in 40 mM Tris, 10 mM MgCl_2_ at pH 8.0 (Tris/Mg^2+^ pH 8.0); and in a typical DON folding buffer (FOB) containing 1 × TAE [40 mM Tris, 19 mM acetic acid, 1 mM ethylenediaminetetraacetic acid (EDTA)] and 12.5 mM MgCl_2_ at pH 8.0. The triangle DON was selected, as it has been widely used as a DOX carrier in previous studies.

The protocol for loading (Figure 3a) was adapted from previous studies (33, 35–39). In the loading process, DONs (here, triangle DONs at 2.5 nM concentration) are mixed with an excess of DOX, the mixture is incubated at RT, and the DOX-loaded DONs are purified from free DOX by *e.g.* centrifugation or spin-filtration. In our experiment, the DOX loading concentration (*c*_0_) was varied from 20 μM to 2 mM. Interestingly, DOX loading has often been performed with DOX concentration in the range of 1–2.5 mM in a FOB containing 10–12.5 mM Mg^2+^ at pH 8.0–8.3. At this pH near the p*K*_a_ of the 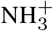 group, part of the DOX molecules are in the deprotonated (uncharged) form and known to be poorly soluble (0.3 mg mL^-1^; 0.55 mM) (47). In addition, dimerization (*K*_a_ = 1.4×10^4^ M^-1^) (45), oligomerization (47), or Mg^2+^ complexation (63–65) can be expected to lead to DOX aggregation.

**Figure 3.**
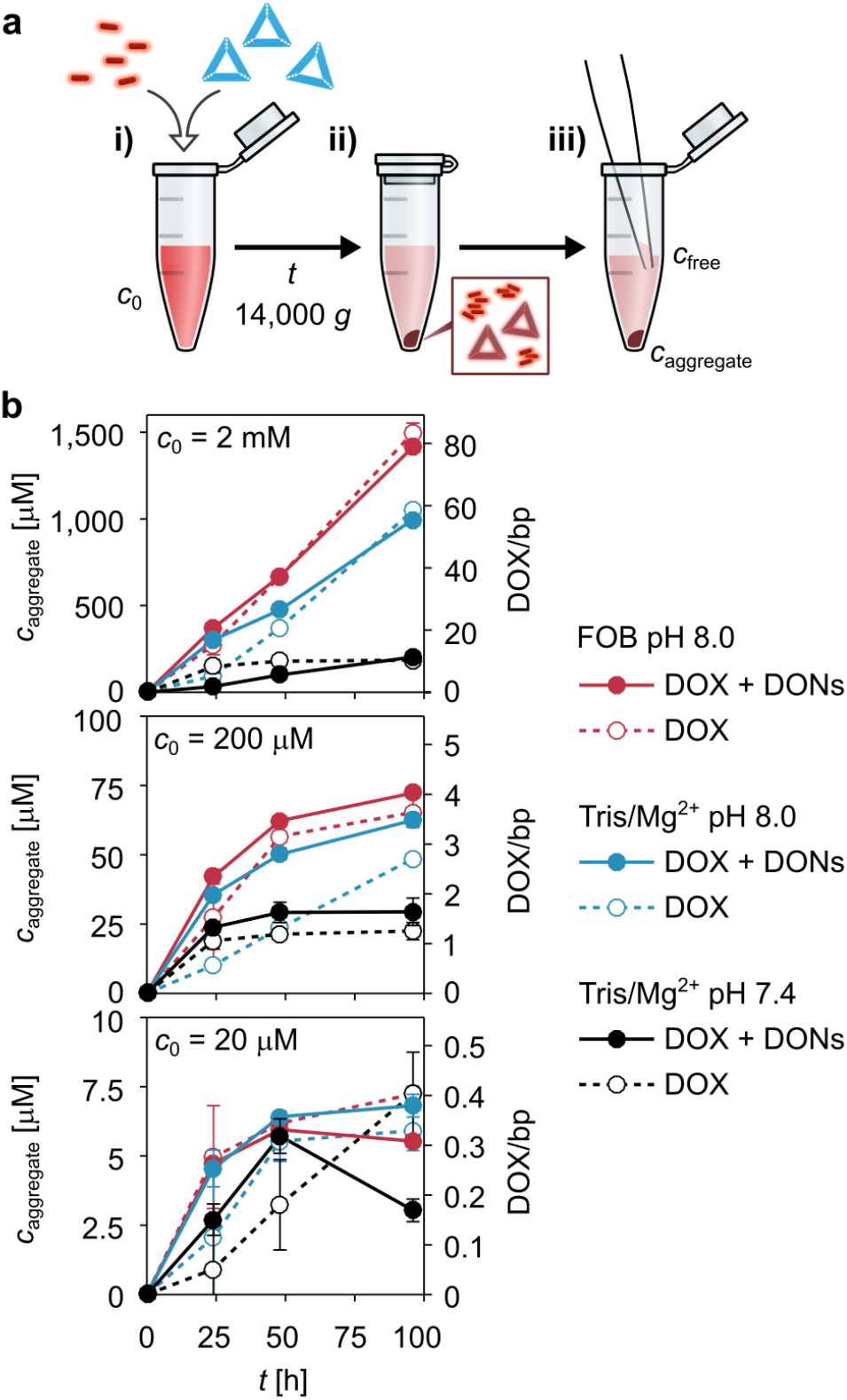
The formation of DOX-DON complexes and DOX aggregates during the loading process. **a**. In the applied loading protocol, i) DOX (*c*_0_ = 2 mM, 200 μM, or 20 μM) is mixed with 2.5 nM triangle DONs and incubated at RT. Control samples are prepared without DONs. ii) After *t* = 24, 48, or 96 h, centrifugation for 10 min at 14,000 *g* is used to separate high-MW particles from the solution; either DOX-loaded DONs or DOX aggregates. iii) The concentration of DOX removed from the solution by precipitation (*c*_aggregate_) is quantified by removing a small volume of the supernatant and determining the DOX concentration in the supernatant (C_free_) from DOX absorbance (A_480_); *C*_aggregate_ = *c*_0_ – *c*_free_. **b**. The concentration of DOX in the pellet (Caggregate) vs. incubation time in the presence and absence of DONs. The amount of sedimentation was determined for 2 mM, 200 μM, and 20 μM DOX loading concentration (*c*_0_) in three different buffers: FOB pH 8.0 (1 2.5 mM MgCl_2_), Tris/Mg^2+^ pH 8.0 (10 mM MgCl_2_), and Tris/Mg^2+^ pH 7.4 (10 mM MgCl_2_). For the DOX-DON samples, the DOX/bp ratio additionally indicates the number of DOX molecules in the precipitate per DNA base pair (*c*_bp_ = 18 μM for 2.5 nM DONs). The *c*_aggregate_ values are expressed as the mean ± standard error, *n* = 3.

During centrifugation, high molecular weight (MW) particles, such as DOX-loaded DONs, are separated from free DOX (low MW) through sedimentation into a dark red precipitate (Figure 3a, middle panel; photographs in the Supplementary Figure 3a). In addition to DOX-loaded DONs, the centrifugation can lead to a sedimentation of other high-MW particles, such as DOX aggregates. To distinquish the DOX-DON formation from the possible aggregation and sedimentation of DOX, each DOX-DON sample in the experiment was compared to a reference sample containing only DOX.

The concentration of DOX in the pellet (*c*_aggregate_) was quantified by determining the concentration of DOX remaining in the supernatant (c_free_) from DOX absorption at 480 nm (Figure 3a, right panel). As shown in Figure 3b, the c_aggregate_ increases with the incubation time, c_0_, pH, and the MgCl_2_ concentration. A comparison of the DOX-DON samples and the DOX-only control samples reveals that nearly identical amounts of DOX are found in the pellets of both DOX-DON samples and DOX-only control samples.

For the DOX-only samples, this supports the hypothesis that DOX self-aggregation takes place during prolonged incubation at RT. Aggregation becomes particularly considerable in samples prepared at pH 8.0 at *c*_0_ = 2 mM, above the solubility limit of deprotonated DOX, confirming that aggregation takes place due to the low solubility of the deprotonated molecules. In addition, the DOX-Mg^2+^ interaction appears to lead to some degree of aggregation in all tested conditions and for all *c*_0_ values.

As the DOX-DON interaction causes only little or no considerable increase in the *c*_aggregate_, it is apparent that the main component in the pellets formed in the DOX-DON samples are likewise DOX aggregates. This becomes even more obvious when considering the amount of DOX in the pellet in relation to the amount of DNA base pairs (bp) in the sample; expressed as the DOX/bp molar ratio in Figure 3b. DOX/bp = 1 can be considered an upper limit of DOX intercalated into DONs, where 100% of the intercalation sites in the DONs are occupied by DOX. For both *c*_0_ = 2 mM and *c*_0_ = 200 μM, the DOX/bp ratio rises above DOX/bp = 1 in less than 24 h. These high DOX/bp ratios observed with triangle DONs indicate a considerable contribution of other aggregation and sedimentation mechanisms, and they are in line with previous reports ranging from DOX/bp = 6.9–19 (12–24 h incubation) (37–39) to as high as DOX/bp = 55 (24 h incubation) (33, 35). We note that this is to our knowledge the first time that this type of DOX-only control samples are presented alongside DOX-DON samples to identify the possible role of DOX aggregation to the sample composition. Our results may thus not only describe the DOX aggregation behavior, but also give a simple explanation for the surprisingly high DOX contents reported in previous studies.

#### Absorption and fluorescence properties of the DOX-DON complexes

We then studied in detail the interaction between DOX and DONs in the selected buffer conditions (40 mM Tris, 10 mM MgCl_2_, pH 7.4). In our experimental setup, a 3 μM solution of DOX is titrated with an increasing concentration of DONs, while maintaining a constant DOX concentration, which causes an increasing fraction of the DOX molecules bind to DNA over the titration. The observed changes in DOX light absorption and fluorescence can be used to extract information about the strength and stoichiometry of the non-covalent binding interaction. To determine whether the DOX loading efficiency into DONs can be tuned with the DON design, we performed the analysis for five structurally distinct DONs (Figure 4a). These include three 2D DONs: the triangle (7), a bowtie (55), and a double-L (55), and two 3D DONs: a capsule (25) and a 24-helix bundle (24HB) (Supplementary Figures 14–16 and Supplementary Table 5). The correct high-yield folding and structural integrity of the DONs were verified with atomic force microscopy (AFM) or transmission electron microscopy (TEM) (Figure 4a).

**Figure 4.**
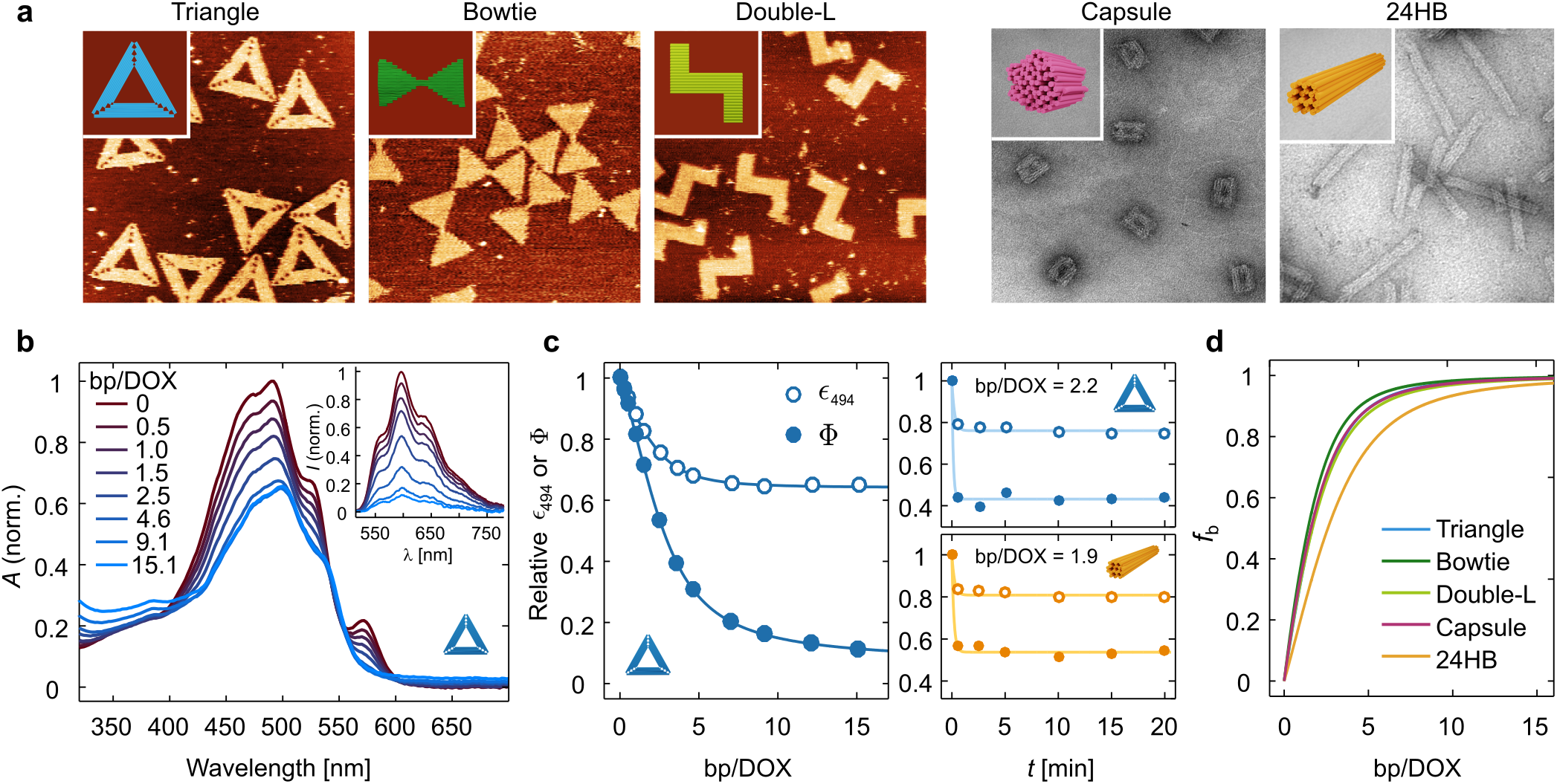
Titration experiments for determining the DOX-loading properties of DONs. **a** The models and microscopy images of the studied 2D and 3D DONs. The triangle, bowtie, and double-L 2D DONs are shown on the left accompanied by atomic force microscopy (AFM) images. The 3D DONs – the capsule and the 24-helix bundle (24HB) are shown on the right in TEM images. The AFM images are 500 nm × 500 nm in size, and the TEM images are 300 nm × 300 nm. **b** Representative changes in the absorption spectrum and fluorescence emission after 494 nm excitation (inset) of 3 μM DOX when the concentration of DNA base pairs (bp) in the solution is increased. The spectra have been measured for the triangle DON after the system has reached an equilibrium. The DNA concentration at each titration step is expressed as the molar ratio between DNA base pairs and DOX molecules in the sample (bp/DOX), and indicated in the legend. The fluorescence spectra have been corrected for the decrease of the molar extinction coefficient at the excitation wavelength (*ϵ*_494_), and represent the quantum yield of the emitting molecules (Φ). **c** The dependency of *ϵ*_494_ and Φ on the bp/DOX ratio (left panel) and the loading kinetics (right panel). In the left panel, the measured values for *ϵ*_494_ and Φ during a titration with the triangle DON have been fitted with a 2-component binding model (Equation 2). The corresponding spectra and titration isotherms for the other DONs are presented in the Supplementary Figure 4. The kinetics of *ϵ*_494_ (empty circles) and Φ (filled circles) in the right panel have been measured by monitoring the absorption and fluorescence spectra of the samples after adding DONs (triangle or 24HB) at the indicated bp/DOX ratio at *t* = 0. The data sets have been fitted with a 1-component exponential decay model of the form *ae*^-bx^ + *c* to illustrate the observed kinetic trends. **d** Increase of the fraction of bound DOX molecules (*f*_b_) when the DNA base pair concentration in the sample increases, obtained by fitting the fluorescence data.

Figure 4b shows the spectral changes of DOX upon titration with the triangle DON. Binding to DNA causes a slight red-shift of the absorption spectrum and an overall decrease of *ϵ* in the visible wavelength region. Additionally, the absorption peak of the DOX-Mg^2+^ complex centered at 576 nm disappears, when the stronger DOX-DNA interaction causes dissociation of the weakly bound DOX-Mg^2+^ complexes. The DOX fluorescence quantum yield (Φ) decreases upon DNA addition, as shown in the inset of Figure 4b for 494 nm excitation. The fluorescence spectra were corrected for the decrease of *ϵ*_494_, which also leads to decreasing fluorescence intensity when less light is absorbed in the sample. The dependency of both *ϵ*_494_ and Φ on the bp/DOX ratio in the sample (titration isotherms) are presented in the left panel of Figure 4c. The results obtained for bowtie, double-L, capsule, and 24HB DONs appear highly similar to the triangle DON, and are presented in the Supplementary Figures 4–5.

**Figure 5.**
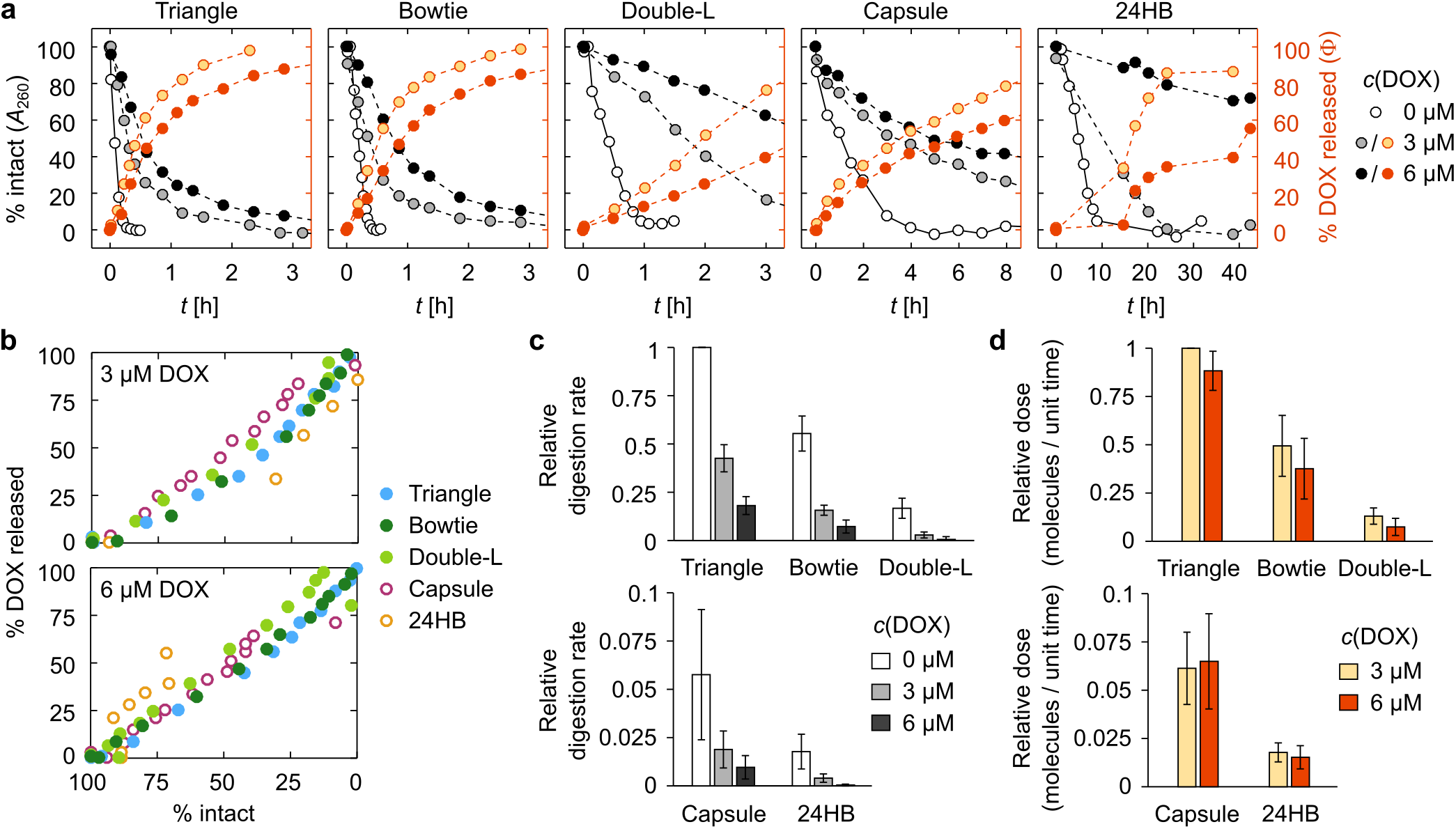
DNase I digestion of the DONs and the subsequent DOX release. **a** Representative (*n* = 1) digestion and DOX release profiles of the studied DONs at different DOX loading concentrations, after introducing 34 U mL^-1^ DNase I into the sample at *t* = 0. The structural integrity of the DONs (% intact) has been determined from the increase of the A_260_ signal, and shown with the white, gray, and black markers depending on the DOX concentration (0, 3, or 6 μM). For samples containing 3or6 μM DOX, the DOX release (% DOX released) is shown with the light and dark orange markers and represents the percentage of initially bound molecules that have been released due to the digestion. % DOX released has been determined from the recovery of the fluorescence quantum yield of DOX. **b** The cross-correlation between % DOX released and % intact for all DONs at 3 and 6 μM DOX loading concentration. **c** DNase I digestion rates of the DONs at 0, 3, or 6 μM DOX loading concentration. All digestion rates are shown relative to the triangle DON at 0 μM DOX concentration, and have been averaged from the digestion rates determined from three individual measurements; expressed as the mean ± standard error. **d** DOX release rates of the DONs at 3 and 6 μM DOX loading concentration. Relative dose stands for the absolute number of DOX molecules released per unit of time. All values are expressed as the mean ± standard error (*n* = 3).

After addition of DNA, the samples were incubated for 2.5 min before the spectra were collected in order to reach an equilibrium where the amount of DOX bound to the DONs has stabilized. The 2.5 min incubation time was found to be sufficient for both 2D and 3D structures, as studied in a separate experiment for both the triangle and the 24HB (Figure 4c, right panel; spectra shown in the Supplementary Figure 8). After addition of DONs, both *ϵ*_494_ and Φ had stabilized by 30 seconds of incubation, and no further spectral changes were observed over longer incubation times. This is in accordance with existing literature where the equilibrium is reached within seconds (45), and shows that the DOX-DON loading process is likewise a faster process than has been previously acknowledged.

Increasing the amount of DONs in the sample causes a discernible scattering effect, which is stronger for the 3D structures than for the 2D structures (Supplementary Figure 10). In the absorbance measurement, this is observed as a slight elevation in the spectrum baseline during the titration. Further analysis of the binding was thus based on the fluorescence data, which is less affected by the light scattering. Analysis of the absorption data was then carried out using the parameters obtained from the analysis of the fluorescence data.

#### Interpretation of the experimental results through a molecular binding model

DOX has been proposed to bind doublestranded DNA (dsDNA) through two prevalent mechanisms: intercalation between G–C base pairs, and minor groove binding at A–T rich areas driven by electrostatic interactions (45). The fluorescence of DOX has been shown to be fully quenched in the strongly bound DOX-GC complex, while the weaker DOX-AT complex remains gently fluorescent (45, 66). For describing the observed decrease of Φ and *ϵ* when increasing the concentration of DONs, we thus applied a 1:2 molecular binding model for including both modes of interaction and the formation of two distinct DOX-DNA complexes with different association constants (*K*_11_ and *K*_12_) and fluorescence quantum yields (Φ_11_ and Φ_12_). This is also fully supported by our observations: when DOX electrostatically binds to a single-stranded DNA (ssDNA), its fluorescence is not quenched, although its absorption spectrum changes in a similar fashion as in the case of DONs (Supplementary Figure 9). However, it is noteworthy that the staple mixture for folding the triangle DON quenches DOX efficiently, as dsDNA residues are formed through partial hybridization or self-complementarity of the staple strands. Therefore, the purification of the structures from the excess staples after DON folding is required for reliable quantification of DOX loading efficiency. As DOX can be assumed to bind selectively in the dsDNA regions of the DONs, the ssDNA nucleotides in the DONs (unpaired scaffold regions and the inert poly-T sequences at the end of the helices) were excluded from the binding analysis.

In the left panel of Figure 4c, the dependence of Φ and *ϵ*_494_ on the base pair concentration (triangle DON) is described according to Equation 2 [Equations (1–7) can be found in the Materials and methods section]. The model suggests that the two DOX-DNA complexes with an average *K*_11_ = (2.0 ± 0.3) × 10^5^ M^-1^ and *K*_12_ = (2.6 ± 0.2) × 10^5^ M^-1^ can be differentiated from each other by the extent of fluorescence quenching (Φ_11_/Φ_0_ = 0.52 ± 0.07 and Φ_12_/Φ_0_ = 0.067 ± 0.009), but in terms of light absorbance their physical properties are similar (*ϵ*_11_/*ϵ*_0_ = 0.58 ± 0.08 and *ϵ*_12_/*ϵ*_0_ = 0.67 ± 0.09 for 494 nm).

While obviously a simplified model of the DOX-DON interaction, the selected binding model can thus be seen to present a reasonable approximation for the behavior of the system and the changes of the physical properties of DOX (Φ and *ϵ*) upon DNA addition by taking into account the two types of binding modes, and essentially, their distinct fluorescence properties. The determined values of *K*_11_ and *K*_12_ are in the same range and order of magnitude as in previous studies (45, 66) – nevertheless, we note that generalization of the fitting results and the obtained parameters outside the presented experimental conditions should be carried out with caution due to the simplifications of the model. A comparison of the fitting parameters for all DONs presented in this study can be found in the Supplementary Table 2.

Finally, the fraction of bound DOX molecules at each bp concentration can be obtained from the fit according to Equation 3, which enables a comparison of the DOX loading properties of the studied DONs (Figure 4d). It appears that the DNA origami superstructure has relatively little effect on how much DOX is bound to the structures, as all curves in Figure 4d are rather similar. In the beginning of the titration, the fraction of bound DOX increases sharply when DNA is introduced into the sample, and the maximum number of bound DOX molecules per base pair is reached at 0.36 ± 0.10 (Supplementary Figure 6).

### DOX release upon nuclease degradation

After determining that all the studied DONs have a similar DOX loading capacity, we studied their differences towards DNase I digestion both in the absence of DOX and when loaded with DOX. For the experiment, DOX-loaded DONs (2 nM) were prepared at both 3 μM and 6 μM DOX loading concentrations. The loading concentration can be used to adjust the DOX loading density. Here, increasing the DOX concentration from 3 μM to 6 μM, while keeping the concentration of DNA base pairs constant, promotes DOX association with the DONs and leads to a higher density of bound DOX molecules. According to the measured quantum yield of DOX and the thermodynamic binding model, the density of bound DOX molecules was 0.17 ± 0.02 DOX/bp for the 3 μM loading concentration and 0.25 ± 0.03 DOX/bp (~47%. higher) for the 6 μM loading concentration. In other words, on average 71% and 63% of DOX molecules in the samples were bound to DONs at 3 μM and 6 μM DOX concentrations, respectively (Supplementary Table 4).

To both confirm the increased loading density at 6 μM DOX concentration and to investigate methods for removing the remaining free DOX from the samples after loading, a supporting experiment was performed: the triangle DON was loaded with DOX at 3–20 μM DOX loading concentration, purified of free DOX with either spin-filtration or with PEG precipitation, and the DOX and DNA contents of the purified samples were measured (Supplementary Figure 12). The determined loading densities were 0.10 ± 0.01 DOX/bp for 3 μM DOX and 0.16 ± 0.02 DOX/bp for 6 μM DOX, with a maximum loading density at 0.45 ± 0.04 DOX/bp. In line with the thermodynamic model, the loading density thus increased by ~60%. when the DOX concentration was doubled. Additionally, both spin-filtration through a 100 kDa MW cut-off membrane and PEG precipitation were termed effective for removing free DOX from DOX-DON samples when the loading was performed at a low (< 10 μM) DOX concentration (purification details are presented in the Supplementary Methods, in the Supplementary Note 13, and in the Supplementary Figures 11–13). However, removing the free DOX disturbs the binding equilibrium, and DOX will be released until a new equilibrium is reached. We importantly observed a fast spontaneous, diffusive release of DOX: freshly spin-filtered, DOX-loaded bowtie DONs had released up to 70% of the initially bound DOX molecules and stabilized into a new binding equilibrium within 10 minutes of the purification (Supplementary Figure 13). The purification of DOX-loaded DNA origami is therefore redundant; while the free DOX concentration is decreased by 80–90%, the DOX loading density of DONs also decreases significantly. The DNase I experiments were thus performed without further purification of the samples.

During DNase I digestion, the nucleases cleave the DONs into short ssDNA fragments. As the *ϵ*_260_/nt for ssDNA is higher than the *ϵ*_260_/nt for dsDNA (67), the process can be followed from the *A*_260_ of the sample. When the DONs are digested, their *A*_260_ increases until reaching a saturation point where the structures are fully digested. The bound DOX is released when the double-helical DNA structure unravels, observed as a recovery of DOX fluorescence. In order to follow both processes in detail during the digestion, we employed a simultaneous kinetic spectroscopic characterization of both the increase of the *A*_260_ and the recovery of DOX fluorescence quantum yield.

Comparing the digestion profiles of the different DONs reveals that the DONs break down at distinct rates depending both on the DNA origami superstructure and the DOX concentration in the sample (Figure 5a). In the beginning of the digestion, the dependence of *A*_260_ on the digestion time is roughly linear. Determining the digestion rates from the linear phase allows a detailed comparison of the DNase I resistance of the different samples (Figure 5c). Both of the studied 3D DONs are digested slower than the 2D structures. The fastest digestion was observed for the triangle DON in the absence of DOX, with the structures being completely degraded within 20 minutes of incubation in the presence of 34 U mL^-1^ DNase I. Loading the DONs with DOX (3 μM) slowed down the digestion considerably, and increasing the loading density further by applying a 6 μM DOX concentration led to a further inhibition of the DNase I activity.

Figure 5a shows how the bound DOX molecules are released from the DONs during the digestion, as determined from the recovery of DOX quantum yield. The release profiles of the studied DONs are in line with the superstructure- and DOX loading concentration dependent trends observed for the structural degradation (Figure 5a). A clear correlation between the fraction of released DOX and the intact dsDNA residues in the sample can be seen for both loading concentrations (Figure 5b). This confirms that the release of DOX is caused purely by the DNase I digestion of the dsDNA framework when the sample is in an equilibrium. The 3 μM DOX samples are digested faster than the 6 μM DOX samples, and the bound DOX is likewise released in a shorter period of time.

Although the 6 μM DOX samples are digested slower than the 3 μM samples, they are also more densely loaded with DOX. This leads to a release of more DOX molecules per a digested DNA base pair. In Figure 5d, the DOX release is thus considered in terms of the relative dose, *i.e.* the number of DOX molecules released per unit of time per unit of DNase I. This decreases the difference between the release properties of 3 μM and 6 μM DOX samples, but does not exceed the effect of the DNase I inhibition. Altogether, the 3 μM samples thus contain less DOX in total and release it faster into the solution, while the loading concentration of 6 μM leads to a slower release of a higher number of loaded DOX molecules.

The spectroscopic results of the DON digestion were also confirmed with an agarose gel electrophoresis (AGE) analysis that was carried out parallel to the spectroscopic experiments (Supplementary Figure 15). The digestion of the structures led to both a lower band intensity and an increased electrophoretic mobility in AGE, correlating well with the % intact values obtained from the spectroscopic measurement. A comparison of the DNase I digestion of unfiltered and spin-filtered bowtie DONs is presented in the Supplementary Figure 14. As the total DOX concentration and the density of bound DOX was drastically reduced through the spin-filtration, the digestion profiles of 3 and 6 μM DOX-loaded bowties started to resemble the digestion of bare origami (0 μM DOX).

## Discussion

### The choice of conditions for DOX loading

In common experimentation with DONs, the buffer of choice is typically a Tris-HCl or a TAE buffer, either supplemented with on average 10–20 mM Mg^2+^. These conditions have been generally found to be appropriate for stabilizing the DONs: the divalent cations effectively screen the electrostatic repulsion between the negative charges of closely packed phosphate backbones, and the typical pH at 8.0–8.3 is in the optimal buffering range of Tris-based buffers. As it is important to retain the structural integrity of DONs throughout experimental procedures, these conditions are also commonly used together with DOX – particularly during loading the DOX into the DONs. Still, the question of whether these conditions can cause unexpected or undesired behavior of DOX, or change its spectroscopic properties in terms of *ϵ* or Φ in a way that can lead to a misinterpretation of spectroscopic observables, has been left almost entirely unaddressed.

Our results show that DONs and DOX have very different optimal environments, and typical DON buffers supplemented with MgCl_2_ at pH 8.0–8.3 are not suited for unequivocal DOX experimentation. In our spectroscopic analysis, we found that when the pH is at or above 8.0 and the MgCl_2_ concentration is at mM range, the environment will lead to DOX heterogeneity either in terms of charge (deprotonation) or formation of DOX-Mg^2+^ complexes (Figure 2a-c). The effects should be carefully considered when interpreting and comparing spectroscopic results obtained in different buffer conditions. For example, it has been stated that the amount of DOX released from the DNA structures increases with decreasing pH (33), but our results strongly suggest that the observed elevation in DOX absorbance and fluorescence may arise from the high absorbance and emission of DOX at low pH instead of from a higher DOX concentration.

In addition to spectral changes, high pH and Mg^2+^ ions can lead to self-aggregation of DOX during prolonged storage times (Figure 3b). In line with previous literature (45, 47, 63, 64), the two major aggregation pathways indicated by our results are the low solubility (0.55 mM) of deprotonated DOX molecules – most clearly observed at pH 8.0 for 2 mM DOX – and the Mg^2+^ complexation, which causes slight DOX aggregation and precipitation regardless of the DOX concentration (from μM to mM range). Importantly, the aggregation of DOX can take place also when preparing DOX-loaded DONs over common incubation times of several hours. The effect of aggregation is most dramatic at poorly optimized conditions (2 mM DOX, pH 8.0, and 12.5 mM MgCl_2_) (Figure 3) and can significantly complicate the purification of free DOX from DOX-loaded DONs from downstream applications. In particular, we observed that upon centrifugation, DOX aggregates precipitate alongside DOX-loaded DONs and can even form the major component of the precipitate. The centrifugation method has been used for purification of both free DOX (33, 35, 37, 40) and daunorubicin (29) from drug-loaded DONs. Our results indicate that the method may lead to a risk of misidentifying aggregated DOX as DOX-DON complexes, and raise questions on the validity of conclusions made about the therapeutic efficiency of the prepared DOX-loaded DONs.

According to our results in the titration experiment, the binding equilibrium between DOX and DONs is reached in less than one minute regardless of the DNA origami superstructure (Figure 4c). An optimal loading time for producing structurally well-defined DOX-loaded DONs thus appears to be in the order of minutes instead of the commonly used incubation times in the range of 12–24 h, and a short loading time may perhaps be the most efficient approach for preventing DOX aggregation during loading. Moreover, we have shown that DOX-loaded DONs prepared at low (< 10 μM) DOX concentrations over short (1 h) loading times can be efficiently purified from free DOX using 100 kDa spin-filtration (34, 36, 38, 39) or PEG precipitation (59) (Supplementary Note 13). However, it is also noteworthy that the sample recovery quickly decreases with an increasing DOX concentration and that the purification may also lead to rapid dissociation of bound DOX (Supplementary Figures 12–13).

### Features of DOX-loaded DONs

In our titration experiments, we studied and compared the DOX-loading capacities of five structurally distinct DONs. The different geometries and design choices of the tested 2D structures (triangle, bowtie and double-L) lead to differences in their flexibility and nuclease accessibility (49), which could cause subtle differences in their DOX loading and release properties. The capsule and 24HB DONs contain a more compact 3D-arrangement of DNA helices. The closed capsule DON is a roughly spherical (31 nm × 28 nm × 33 nm) structure with a hollow inner cavity (25). Such hollow architectures can be expected to make DNA intercalation sites more accessible to DOX, and therefore they have been previously applied for enhancing the loading efficiency of daunorubicin (29). On the other hand, the DNA helices in the 24HB are arranged into a tight, regular bundle of ~12 nm in diameter and 115 nm in length.

It is thus interesting to note that in our experimental conditions and in the titration experiment presented in Figure 4, roughly identical amounts of DOX were incorporated into all of the tested DONs in terms of density of the drug molecules in the DONs. The maximum DOX loading content was determined to be 0.36 ± 0.10 DOX molecules bound per one DNA base pair. The value is in line with a number of previous studies for calf thymus dsDNA, where the maximum binding efficiency of DOX had been determined to be in the range of ~0.29–0.37 DOX molecules per base pair (45, 68). The similarity of the tested DONs is a rather surprising observation, as the steric hindrance from the compact arrangement of DNA helices could be expected to lead to a restricted accessibility of DNA helices and intercalation sites particularly in the 3D DONs. In fact, such kind of restricted loading has recently been observed for the bis-intercalator YOYO-1 (70). The different behavior we observe for DOX might arise from the different binding mechanisms of the two drugs, with bis-intercalation being more affected by the steric hindrance.

Our observations of a low DOX-loading capacity of DONs are additionally contradictory to many previous studies on DON-based DOX delivery, where the reported concentrations of bound drug molecules in DONs are often higher than the concentration of DNA base pairs in the sample – up to 113 DOX molecules per base pair (33). While it is rather obvious that intercalation cannot be the only DOX binding mechanism behind the previous reported high DOX loading contents, the other possible processes, such as minor-groove binding or even external aggregate formation through stacking interactions (45), are rarely discussed. Our results support the interpretation that all three mechanisms might play a role in the loading process depending on the choice of experimental conditions. An atomic force microscopy (AFM) – based characterization of DOX-loaded double-L DONs (Supplementary Figure 7 and Supplementary Table 3) shows that the DOX binding causes a torsional twist in a concentrationdependent manner, indicative of an intercalative binding mode (34, 69). Our spectroscopic results also strongly support the presence of a minor-groove binding mode that leads to a lesser extent of DOX fluorescence quenching (45, 66). In addition, stacking into aggregates on the DON surface or the observed self-aggregation of DOX could present a plausible explanation for the previous DOX loading contents well above a loading density of DOX/bp > 1, and result from suboptimal buffer conditions or prolonged incubation times during loading.

In addition to the kinetics of the loading process, a second crucial kinetic parameter of DOX-loaded DONs is the rate of diffusive release of DOX in low-DOX conditions. Interestingly, we observed rapid spontaneous DOX release after the DOX-loaded structures were purified from excess DOX. As seen in the data presented in Supplementary Figure 13, freshly spin-filtered DOX-loaded bowtie DON samples had reached a new equilibrium state through DOX release (recovery of the quantum yield) in less than 10 minutes. No further quantum yield increase was observed between 10 minutes and 45 hours. To our knowledge, such rapid diffusive release has not been previously reported for DOX-loaded DONs, albeit it is in line with the relatively large dissociation rate constant (*k*_off_) of DOX. Different sources have reported the *k*_off_ of DOX as 2.07–8.49 s^-1^ at 25 °C (71, 72), giving an equilibration half-time of *t*_1/2_ = ln2/*k*_off_ = 0.08–0.33 s. From the pragmatic viewpoint, this indicates that although the amount of free DOX in the solution can be reduced through purification, it simultaneously decreases the amount of bound DOX in DONs, and it seems impossible to eliminate free DOX entirely (decrease of free DOX leads to dissociation of bound DOX). The rate of diffusive release likely depends on the sample composition and the DOX binding mode – intercalation, external binding, or aggregation – and the implications for downstream drug delivery applications would in turn present a subject for further studies.

### DNase I digestion leads to DOX release at superstructure-dependent rates

As an important prerequisite for biomedical applications, we simulated the possible degradation pathways of the complexes in nuclease-rich environments. DNase I was selected as a degradation agent, as it is the most important and active nuclease in blood serum and mammalian cells. In addition, DON digestion by DNase I has been previously studied (49, 51, 57, 73, 74), but not with this kind of an approach that allows detailed simultaneous monitoring of DON cleavage and drug release.

The superstructure-dependent DON degradation rates were resolved by following the increase of DNA absorbance at 260 nm. The stability varies from structure to structure, which has also been observed in the previous studies. In general, the DNase I digestion is notably slower for 3D than 2D structures; in the most extreme case (triangle vs. 24HB), by roughly two orders of magnitude (Figure 5a and 5c). The plain 2D structures contain flexible regions that are likely more susceptible to DNase I digestion (75), and they follow similar digestion profiles as reported earlier (49). The increased stability of the 24HB compared to the capsule may originate from design features such as a more compact, rigid and regular structure and a higher amount of scaffold crossovers, which are all factors known to increase the resiliency of DONs towards nuclease digestion (49–51, 57, 74).

As the percentage of released DOX correlates well with the degradation level of the DONs (Figure 5b), the different digestion rates enable customized drug release over a rather wide time window. When the structures are loaded with DOX, the digestion slows down with increasing DOX concentration and adds one more controllable parameter to the tunable release profile: 6 μM DOX loading concentration yields lower relative doses (released amount of DOX / unit time) than 3 μM (Figure 5d). As the 6 μM concentration leads to increasing density of DOX molecules loaded into the DONs, the underlying mechanism is most likely DOX-induced DNA protection through interference with the minor-groove binding of DNase I. The inhibitory effect of DOX has been previously observed for dsDNA with variable sequences (76), and shown to be caused specifically by the DNA-bound DOX rather than interactions between free DOX and DNase I (77).

Furthermore, to achieve reasonable time scales for the digestion rate comparison, we have here applied a DNase I concentration (34 U mL^-1^) that is remarkably higher than for example in blood plasma (0.36 ± 0.20 U mL^-1^ (78)). As the concentration of DNase I is essentially defined through its activity – affected by *e.g.* the temperature and the salt concentration – the acquired results set an appropriate starting point to estimate the relative susceptibility and the drug release capacity of distinct DNA shapes in DNase I-rich environments. Obviously, DNase I digestion is only one of the factors compromising the stability of DONs in a physiological environment (53, 54). A more complete picture of the durability of DONs in DOX delivery, i.e. the combined effects of the physiological cation concentrations, temperature, and interactions with other plasma proteins than DNase I presents an interesting topic for future studies. Here, by studying the DNase I digestion of distinct DONs in isolation from the other destabilizing factors, we have been able to resolve in detail their stability as DOX carriers and their superstructure-dependent differences related to their resistance against nucleases. In a nutshell, the various DON shapes used in this work and the applied DOX-loading levels together provide a broad selection of relative doses for engineered DOX delivery (Figure 5d).

## Conclusions

We have shown that the release of the common therapeutic drug DOX from DONs upon DNase I digestion can be customized by rationally designing the DNA superstructures and adjusting the loading concentration of DOX. In our model system, we observed clear correlation between the released DOX and the degradation level of the DONs. Both the superstructure and rigidity of DONs have an impact on their stability against nucleases, which is in agreement with previous studies (49, 51, 57). The stiffness and resilience of DONs achieved by the close packing of helices may, on the other hand, deteriorate the loading capacity of DNA-binding drugs (70). Nevertheless, here we observed nearly identical DOX loading properties for all tested DONs in terms of the loading time and the loading yield, but drastically different digestion and release profiles. Increasing the amount of loaded DOX slows down the digestion, which is plausibly associated with restricted DNase I cleavage due to the interfering DNA-bound DOX (76, 77).

Importantly, our spectroscopic analysis of free DOX, the DOX-DON loading process, and the DOX-loaded DONs under different conditions reveals that a number of studies have inadequately estimated the DOX loading capacity of DONs and overlooked the potential effect of DOX self-aggregation during sample preparation and experimentation. In addition, we present previously unacknowledged fast kinetics of DOX loading and spontaneous release after removal of the free DOX. We propose that suboptimal conditions and procedures in sample preparation and characterization may lead to vaguely defined sample composition and to misleading interpretation of the actual drug efficacy. Therefore, our results may also help in explaining previous, often incoherent reports on DON-mediated DOX delivery. In addition, to resolve the exact amount of DOX that is bound to DONs through the loading procedure is also a matter of the utmost importance when cost-effectiveness of DON-based targeted delivery is assessed (21).

Our observations underline the significant potential of DONs in drug delivery applications and provide guidelines for choosing appropriate protocols for preparing, studying, purifying, and storing DOX-loaded DONs. Here we employed plain DONs without further modifications, but by taking advantage of their unsurpassable addressibility and modularity, multifunctionalities can be further realized. In the bigger picture, we believe our findings will help in building a solid ground for the development of safe and more efficient DNA nanostructure-based therapeutics with promising programmable features.

## Data availability

The authors confirm that the data supporting the findings of this study are available within the article and the Supplementary Information files. Some parts of the raw data are available from the corresponding author, upon reasonable request.

## Supplementary data

Supplementary Information accompanies this paper.

## Author contributions

H.I. performed the majority of the experiments, modeled the data and wrote the manuscript. B.S. and A.H.-J. performed additional experiments. A.K. conceived the project. M.A.K., T.L. and J.A.I. supervised the work. V.L. conceived, initialized and supervised the project, and wrote the manuscript. All authors designed the experiments, analyzed and discussed the data, and also commented on and edited the manuscript.

## Acknowledgements

We thank Dr. H. Häkkänen for technical assistance and S. Julin for the 24HB DNA origami design. We acknowledge the provision of facilities and technical support by Aalto University Bioeconomy Facilities and OtaNano – Nanomicroscopy Center (Aalto-NMC). The research was carried out under the Academy of Finland Centres of Excellence Programme (2014–2019).

