## Supplementary Information for "Unraveling the interaction between doxorubicin and DNA origami nanostructures for customizable chemotherapeutic drug release"

### Supplementary methods

#### Materials

All DOX solutions were prepared by diluting 10 mM stock solution of DOX (in deionized (DI) water) with the applied buffer. When preparing solutions with high DOX concentrations ($\geq$ 200 μM), DOX was diluted with DI water and 2× buffer at 1/2 of total sample volume to prevent buffer dilution.

The abbreviations of the buffers applied in the experiments are: Tris/$Mg^{2+}$ pH 7.4 = 40 mM Tris, 10 mM $MgCl_2$, pH 7.4; Tris/$Mg^{2+}$ pH 8.0 = 40 mM Tris, 10 mM $MgCl_2$, pH 8.0; FOB pH 8.0 = 1× DON folding buffer: 1× TAE [40 mM Tris, 19 mM acetic acid, 1 mM ethylenediaminetetraacetic acid (EDTA)] and 12.5 mM $MgCl_2$, pH 8.0.

The spin-filtration for removing free DOX from DOX-DON samples was carried out using Amicon Ultra 0.5 mL centrifugal filter units (Merck Millipore) with 10 kDa, 30 kDa, or 100 kDa molecular weight cutoff.

#### Additional DOX self-aggregation experiments

The results presented for DOX self-aggregation in Figure 3 in the main article were complemented with supplementary experiments of DOX self-aggregation. To study whether the chosen centrifugation procedure (10 min centrifugation at 14,000 $g$) for separating free DOX from high molecular weight particles can affect the obtained $c_{\text{aggregate}}$ values, the experiment was repeated for 200 μM DOX samples using different centrifugation speeds. In addition, the self-aggregation of 3 μM DOX in the Tris/$Mg^{2+}$ pH 7.4 buffer was studied over 96 h incubation.

**The effect of the centrifugal force on the $c_{\text{aggregate}}$.** DOX solution at 200 μM concentration was prepared in FOB pH 8.0. To obtain a $t$ = 0 value for $A_{480}$ and the $c_{\text{aggregate}}$, UV-Vis spectrum of the sample between 240–600 nm was collected from a 2 μL droplet on a BioTek Take3 plate with a 0.05 cm optical path using a BioTek Eon microplate spectrophotometer. The DOX solution was then divided into four aliquots, which were incubated for 24 h at RT. After the incubation period, the aliquots were centrifuged for 10 min at either 2,000, 6,000, 10,000, or 14,000 $g$, small volumes of the supernatants were removed, and the UV-Vis absorption spectra were recorded similarly to the $t$ = 0 spectra. The $c_{\text{aggregate}}$ values at the 24 h time point were calculated as $c_{\text{aggregate}}$ = 200 μM × $A_{480}(t = 24\text{ h})$ / $A_{480}(t = 0)$. The experiment was repeated three times and the $c_{\text{aggregate}}$ values were reported as the mean ± standard deviation of the three experiments.

**Self-aggregation of 3 μM DOX.** 3 μM DOX solutions were prepared in Tris/$Mg^{2+}$ pH 7.4 and in DI water. The UV-Vis absorbance spectra of the solutions between 240–650 nm were measured directly after preparing the solutions to obtain the $t$ = 0 values for DOX absorbance and concentration in the samples. Spectroscopic measurements were performed with a Varian Cary UV-Vis spectrophotometer for 750 μL of sample in a disposable BRAND micro UV cuvette. Each sample was then aliquoted into three parts and incubated at RT. After 24, 48, or 96 h, one aliquot was centrifuged at 14,000 $g$ for 10 minutes and the UV-Vis spectrum of the supernatant was collected identically to the $t$ = 0 measurement. The $c_{\text{aggregate}}$ values at the different time points were calculated as $c_{\text{aggregate}}$ = 3 μM × $A_{480}(t)$ / $A_{480}(t = 0)$. The experiment was repeated three times and the $c_{\text{aggregate}}$ values were reported as the mean ± standard deviation of the three experiments.

#### Purification of free DOX from DOX-DON samples after loading

**Comparison of DOX retention by 10, 30, and 100 kDa molecular weight cutoff filter membranes.** Spin-filtration through 10 kDa, 30 kDa, and 100 kDa molecular weight cutoff membranes was used both to study the size of DOX aggregates formed in samples containing 2 mM DOX in FOB pH 8.0, and to evaluate the effectiveness of different filter membranes in the purification of free DOX and DOX aggregates from DON-DOX samples after loading. Prior use, the filters were rinsed with the sample buffer (either FOB pH 8.0 or Tris/$Mg^{2+}$ pH 7.4 depending on the experiment) by adding 500 μL of buffer in the filter unit, centrifuging for 10 min at 14,000 $g$, and removing the buffer remaining in the filter with a pipette.

For the comparison of 10 kDa, 30 kDa, and 100 kDa filter membranes, 2 mM DOX solution was prepared in FOB pH 8.0 and incubated overnight at RT to induce formation of DOX aggregates. 100 μL of the DOX solution was pipetted into pre-rinsed 10, 30, and 100 kDa filters and 10 μL of each solution was removed for UV-Vis measurements. The samples were then centrifuged for 5 min at 14,000 $g$. UV-Vis spectra between 240–600 nm were measured from the unfiltered samples, from the filtrates, and from the concentrates (sample remaining in the filter after centrifugation). $c_0$, $c$(filtrate), and $c$(concentrate) values were determined from the $A_{480}$ values. The experiment was repeated total three times and the relative concentrations of DOX were reported as the mean ± standard deviation.

The extent of DOX binding to the 100 kDa filter membrane was further studied by comparing the $c_0$ and $c$(filtrate) values of various DOX-only and DOX-DON samples. DOX-DON samples were prepared by mixing DOX at either 3, 6, 20, or 200 μM concentration with the triangle DON at 2 nM final concentration in Tris/$Mg^{2+}$ pH 7.4, and incubating the samples at RT for 1 h to load the DONs with DOX. DOX-only samples were prepared identically but without DONs. After the incubation, the solutions were pipetted into pre-rinsed 100 kDa cutoff filters and spin-filtered briefly (4 min at 6,000 $g$). UV-Vis absorption spectra were measured both from unfiltered samples and from the collected filtrates. 3 μM and 6 μM samples were analyzed with a Varian Cary UV-Vis spectrophotometer from 750 μL of sample volume in a disposable BRAND micro UV cuvette ($l$ = 1 cm) and UV-Vis spectra were collected between 240–650 nm. Samples containing 20 μM or 200 μM DOX were analyzed with a BioTek Eon microplate spectrophotometer, and the UV-Vis spectra were collected between 240–600 nm from a 2 μL droplet of sample on a BioTek Take3 plate ($l$ = 0.05 cm). Because binding to DONs changes the shape of the absorption spectrum of DOX, the concentration values $c_0$ and $c$(filtrate) were calculated from $A_{544}$ values – *i.e.* absorbance at the isosbestic point. The $c$(filtrate) / $c_0$ values were reported from a single measurement.

**Free DOX removal by spin-filtration.** Spin-filtration for removing unbound DOX from DOX-DON samples after loading was carried out with 100 kDa molecular weight cutoff spin-filters. The buffer used in all samples and purification experiments was Tris/$Mg^{2+}$ at pH 7.4. All incubation and spin-filtration steps of the experiment were performed at RT.

For studying the effect of DOX loading concentration on the loading and purification outcome, DOX-loaded DON samples were prepared at 3, 6, and 20 μM DOX concentrations with all samples containing 2 nM triangle DON. The samples were incubated for 1 h for loading the DONs with DOX. UV-Vis spectra of the samples were measured after the incubation with a Varian Cary UV-Vis spectrophotometer. The samples were then purified from unbound DOX with the following spin-filtration protocol:

1. The filter was rinsed by adding 500 μL of buffer in the filter, centrifuging for 10 min at 6,000 $g$, and discarding the flow-through. The $\sim$20 μL volume of buffer left in the filter was not removed.

2. 480 μL (of total 800 μL) of DOX-DON sample was added, centrifuged for 10 min at 6,000 $g$, and the flow-through was discarded.

3. The remaining 320 μL of sample was added to the $\sim$20 μL of concentrated sample in the filter together with 160 μL of buffer, and centrifuged for 10 min at 6,000 $g$. The flow-through was discarded.

4. 480 μL of buffer was added and the sample was centrifuged for 10 min at 6,000 $g$.

5. The concentrated sample was collected into a fresh microcentrifuge tube by inverting the filter unit and centrifuging for 2 min at 1,000 $g$. The sample was then diluted with 780 μL of buffer to a total volume of 800 μL (original sample volume).

UV-Vis spectra of the purified samples were measured identically as before purification. For comparing the sample composition before and after purification, the concentrations of DOX [$c$(DOX)] and DNA base pairs [$c$(bp)] were quantified from the UV-Vis spectra. $c$(DOX) for each sample was determined from the $A_{544}$ value (isosbestic point) based on a standard curve of $A_{544}$ *vs.* $c$(DOX) for unpurified samples. The $c$(bp) values were calculated from $A_{260}$ after subtracting the contribution of DOX absorbance [$A_{260}$(DOX)] from the measured $A_{260}$: $A_{260}$(DNA) = $A_{260} - A_{260}$(DOX). $A_{260}$(DOX) was estimated as $A_{260}$(DOX) = $A_{544}\times$ [$A_{260}$(DOX, free)/$A_{544}$(DOX, free)], where the ratio of absorbances $A_{260}$(DOX, free)/$A_{544}$(DOX, free) was collected and averaged from DOX-only samples at 3, 6, or 20 μM concentration. DOX/bp ratios were calculated from the $c$(DOX) and $c$(bp) values of the purified samples. The DNA recovery yield was defined as % DNA recovery = $c$(bp) / $c$(bp)$_0 \times 100\%$, where $c$(bp) refers to DNA base pair concentration in the sample after purification, and $c$(bp)$_0$ to the base pair concentration in the unpurified sample. The DOX/bp ratios and the DNA recovery yields were reported as an average ± standard deviation after repeating the experiment three times.

The efficiency of spin-filtration in free DOX removal was measured with DOX-only reference samples: 3, 6, and 20 μM solutions of DOX were prepared in the Tris/$Mg^{2+}$ buffer at pH 7.4, incubated for 1 h at RT, and processed with the same spin-filtration procedure as the DOX-DON samples. The remaining concentration of free DOX in the samples ($c$) was quantified from the $A_{544}$ values. The DOX removal efficiency for each loading concentration was calculated from the difference of $c$ and $c_0$: % free DOX removed = $(c_0 - c)/c_0 \times 100\%$.

**Free DOX removal by PEG precipitation.** In addition to spin-filtration, the efficiency of PEG precipitation as a purification method for removing unbound DOX from DOX-loaded DONs was evaluated. The purification was carried out for an identical set of DOX-DON samples (2 nM triangle DON + 3, 6, or 20 μM DOX) and DOX-only references as for the 100 kDa spin-filtration.

The PEG precipitation was carried out by mixing 750 μL of DOX-DON or DOX-only sample with 750 μL of PEG precipitation buffer (1$\times$ TAE, 505 mM NaCl, 15% (w/v) PEG8000), centrifuging for 30 min at 14,000 $g$, removing the supernatants, and resuspending the pellets to 750 μL of buffer (original sample volume). For DOX-DON samples prepared with 3 μM or 6 μM DOX loading concentration, the pellets could be redissolved by first mixing the samples gently with a pipette followed by overnight incubation at RT. The pellets formed in the DOX-DON samples with 20 μM DOX loading concentration required 24 h incubation under shaking (600 rpm) at +30 °C before being completely dissolved.

Free DOX removal efficiency, DNA recovery, and DOX/bp ratios in the purified samples were determined from UV-Vis absorption spectra with an identical protocol to the spin-filtration analysis.

**Spin-filtration and DNase I digestion of bowtie DONs.** The DNase I digestion of DONs after free DOX removal with spin-filtration, as well as the diffusive release of DOX form DONs after purification was studied with the bowtie DON. The DONs ($\sim$2 nM) were briefly incubated (30–60 min) with 3 or 6 μM DOX to prepare DOX-loaded bowtie DONs. Unbound DOX was then removed with spin-filtration according to the protocol described under the Supplementary Methods subsection "Free DOX removal by spin-filtration", but with the exception that the original sample volume of 921 μL was concentrated to 760 μL in the final buffer addition step to counterbalance for the sample loss in the spin-filtration. The DNase I digestion and DOX release experiment was then carried out as described in the Materials and Methods section of the main text for both the filtered samples and unfiltered DOX-DON samples. The unfiltered samples were diluted before the digestion to a DNA concentration to the concentration of the filtered samples.

The same spin-filtration and DNase I digestion experiment was also carried out for bowtie DONs with 0 μM DOX as a control to ensure that the spin-filtration of plain DONs does not affect the digestion rate. A purification efficiency of $\sim$100% was confirmed by collecting the absorption and fluorescence spectra of 3 μM and 6 μM DOX-only samples before and after a spin-filtration. For studying the diffusion rate of DOX from spin-filtered samples, one set of 3 and 6 μM spin-filtered bowtie DON samples were incubated at RT after the spin-filtration and their absorption and fluorescence spectra were monitored over a 46 hour time period.

### Agarose gel electrophoresis (AGE)

2% agarose gels were prepared in a buffer containing $1\times$ TAE and 11 mM $MgCl_2$ and pre-stained with ethidium bromide (0.47 μg $mL^{-1}$ final concentration in the gel). DON samples were loaded on the gel with loading dye (Sigma-Aldrich). The gel was run for 45 minutes at 90 V on an ice bath, and imaged under UV light with a BioRad ChemiDoc MP imaging system.

### Supplementary notes

#### Note S1: DOX fluorescence in different pH buffers

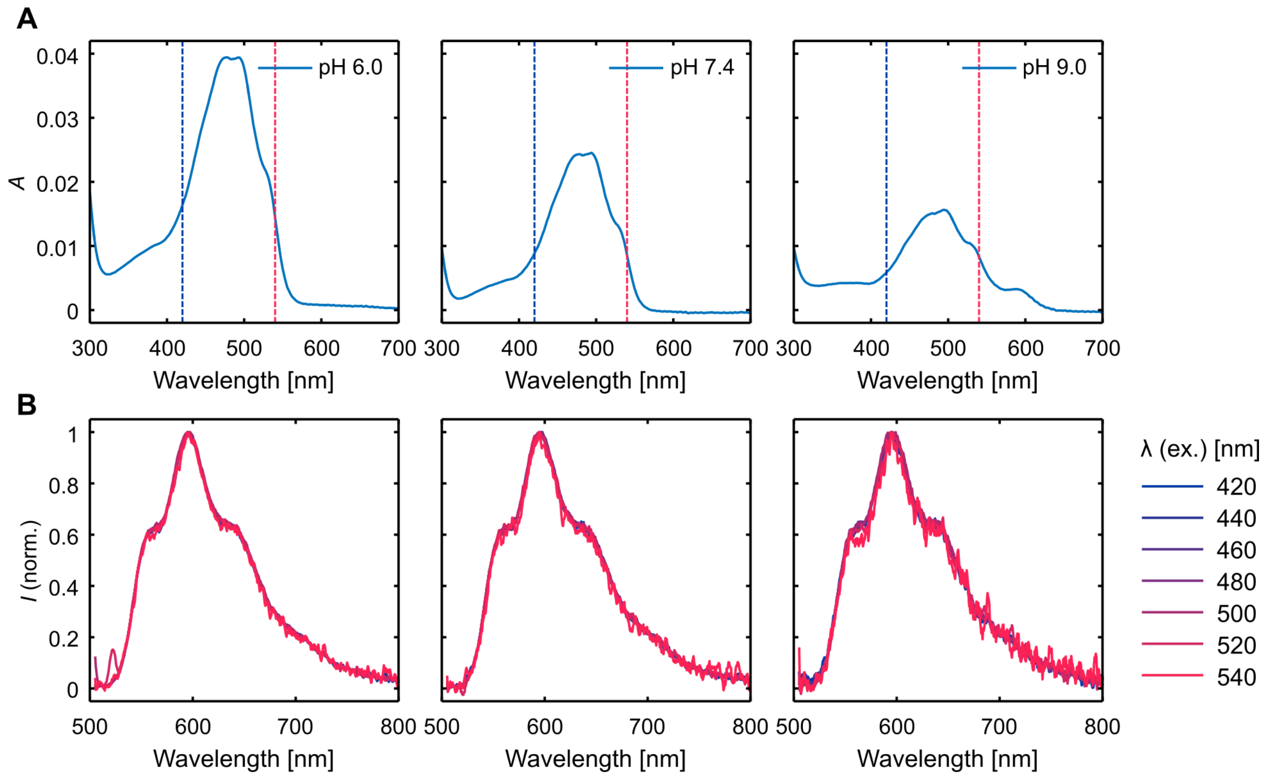

**Supplementary Figure 1.** Comparison of the shape of the absorption and emission spectra of 3 μM DOX in 40 mM Tris-HCl buffer at pH 6.0, 7.4, and 9.0 prepared without magnesium. (**A**) Absorption spectra at pH 6.0, 7.4, and 9.0. The colored dashed lines indicate the excitation wavelength range applied in Figure B. (**B**) The shape of the emission spectra collected with excitation wavelengths in the range of 420–540 nm. All spectra are normalized to the maximum value. While the emission intensity of DOX depends on the excitation wavelength and the pH of the sample (lower emission intensity at higher pH), the shape of the emission spectrum does not change, indicating that the full emission originates from a homogeneous group of fluorescent molecules.

### Note S2: DOX fluorescence spectrum heterogeneity in the presence of different concentrations of $MgCl_2$

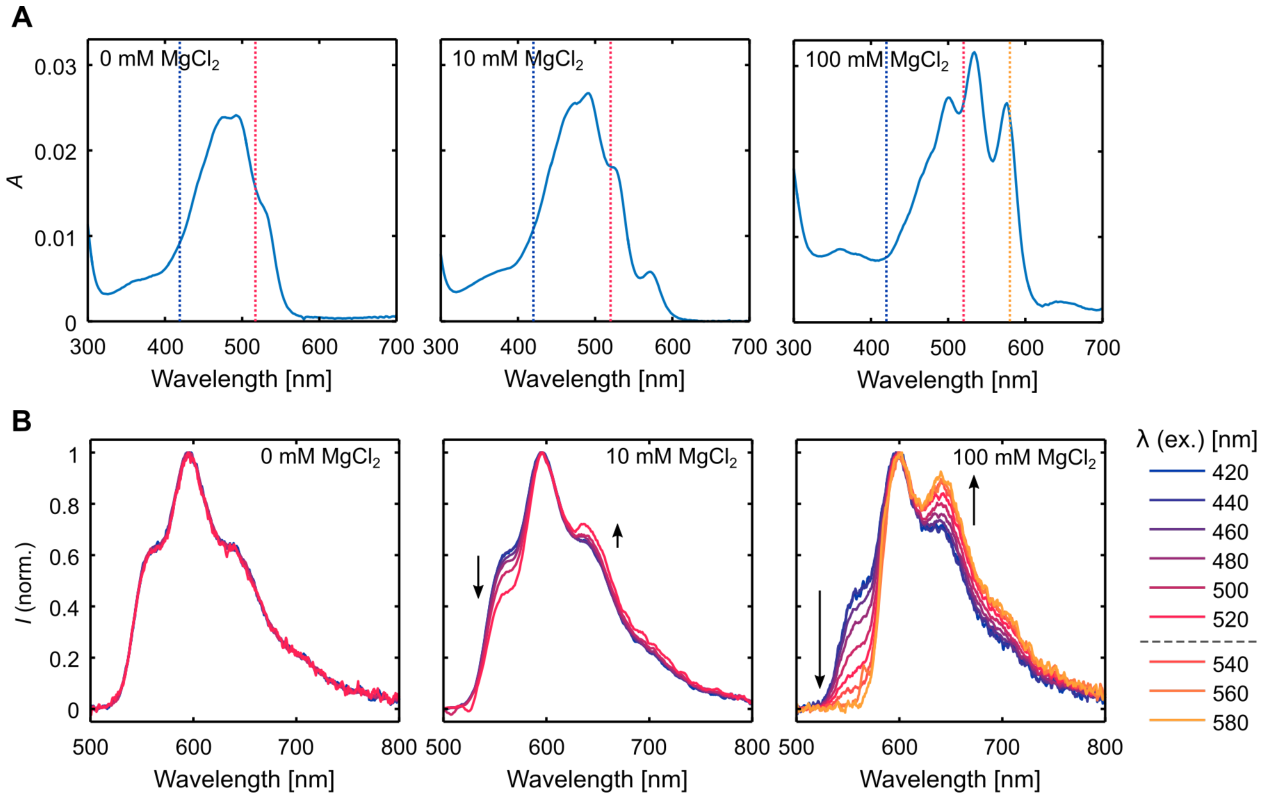

**Supplementary Figure 2.** Absorption and emission spectra of 3 $\mu$M DOX in 40 mM Tris pH 7.4 supplemented with different concentrations of $MgCl_2$. (**A**) Absorption spectra measured at 0, 10, or 100 mM $MgCl_2$ concentration (the concentration indicated in the upper-left corner of each figure). The excitation wavelength range applied for the emission spectra in Figure B (420–520 nm, and 520–580 nm) are indicated with the colored dashed lines. (**B**) Heterogeneity of the emission spectrum of DOX caused by $Mg^{2+}$ complexation. The arrows indicate the changes observed in the shape of the spectrum relative to the spectrum collected with 420 nm excitation when the excitation wavelength changes. For 0 mM and 10 mM $MgCl_2$ samples, the shape of the emission spectrum is compared between 420–520 nm excitation. For the 100 mM $MgCl_2$ sample, additional excitation wavelengths 540–580 are included. The emission spectrum collected with 580 nm excitation originates purely from the DOX-$Mg^{2+}$ complexes, as pure DOX (0 mM $MgCl_2$ sample) does not absorb light at this wavelength. All spectra have been normalized to the intensity at emission maximum.

### Note S3: Supplementary DOX self-aggregation data

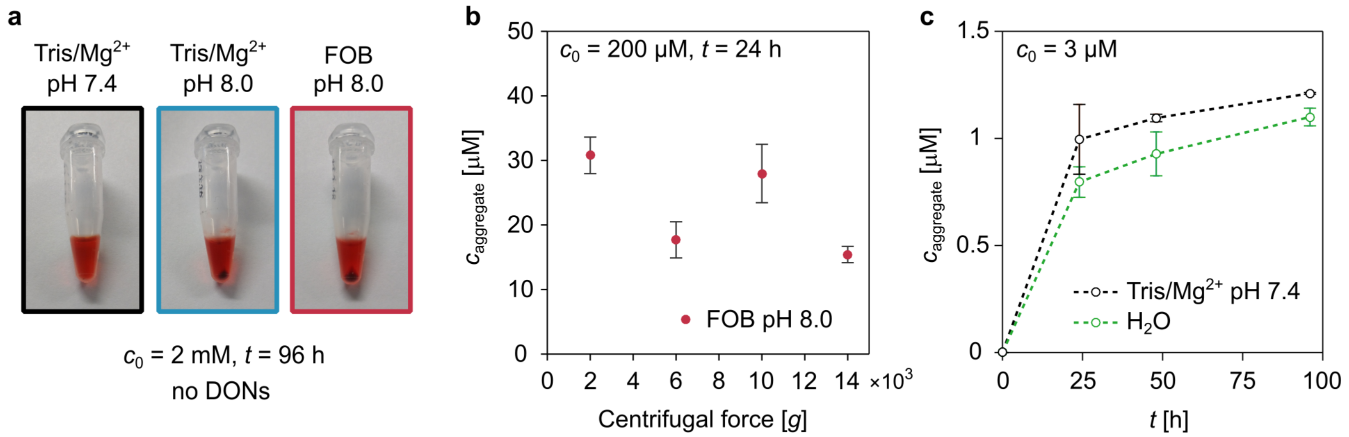

**Supplementary Figure 3.** Supplementary characterization of DOX self-aggregation. (**a**) Photographs of DOX sedimentation in the DOX-DON loading and aggregation experiment (Figure 3 in the main text). DOX solutions ($c_0$ = 2 mM) in the absence of DONs were incubated at RT for 96 hours, followed by centrifugation at 14,000 *g* for 10 min to separate DOX aggregates from the solution. DOX self-aggregation was visible to the eye also for 200 μM DOX samples in both Tris/$Mg^{2+}$ pH 8.0 and FOB pH 8.0, but not for smaller (20 and 3 μM) DOX concentrations or any DOX concentrations in Tris/$Mg^{2+}$ pH 7.4 buffer. (**b**) The effect of centrifugal force on the measured $c_{aggregate}$ in the experiment presented in Figure 3 for 200 μM DOX in FOB pH 8.0 after 24 h incubation at RT. (**c**) Self-aggregation of 3 μM DOX in the Tris/$Mg^{2+}$ pH 7.4 buffer and in DI water.

### Note S4: Details of the DNA origami designs

**Supplementary Table 1.** Nucleotide amounts in the studied DNA origami designs ($N_{\mathrm{total}}$) and the fractions of unpaired ($N_{\mathrm{ss}}$) and hybridized ($N_{\mathrm{ds}}$) nucleotides. The $N_{\mathrm{ds}}$ includes both the base pairs formed between the staple strands and the scaffold, and the base pairs formed as secondary structures in unpaired scaffold regions (simulated with the NUPACK web application). Molar extinction coefficients per nucleotide at 260 nm ($\epsilon_{260}$/nt) have been calculated for each shape according to Equation (1).

| Origami shape | $N_{\mathrm{total}}$ | $N_{\mathrm{ds}}$ | $N_{\mathrm{ss}}$ | $N_{\mathrm{ds}}/N_{\mathrm{total}}$ (%) | $\epsilon_{260}$/nt ($\mathrm{cm}^{-1}\mathrm{M}^{-1}$) |
|---|---|---|---|---|---|
| Triangle | 14,516 | 14,464 | 52 | 99.6 | 6,700 |
| Bowtie | 15,039 | 13,948 | 1,091 | 92.7 | 6,900 |
| Double-L | 15,193 | 14,324 | 869 | 94.3 | 6,900 |
| 24HB | 15,504 | 15,120 | 384 | 97.5 | 6,800 |
| Capsule | 16,732 | 14,704 | 2,028 | 87.9 | 7,000 |

### Note S5: Spectra and titration curves for the bowtie, double-L, 24HB, and capsule origami

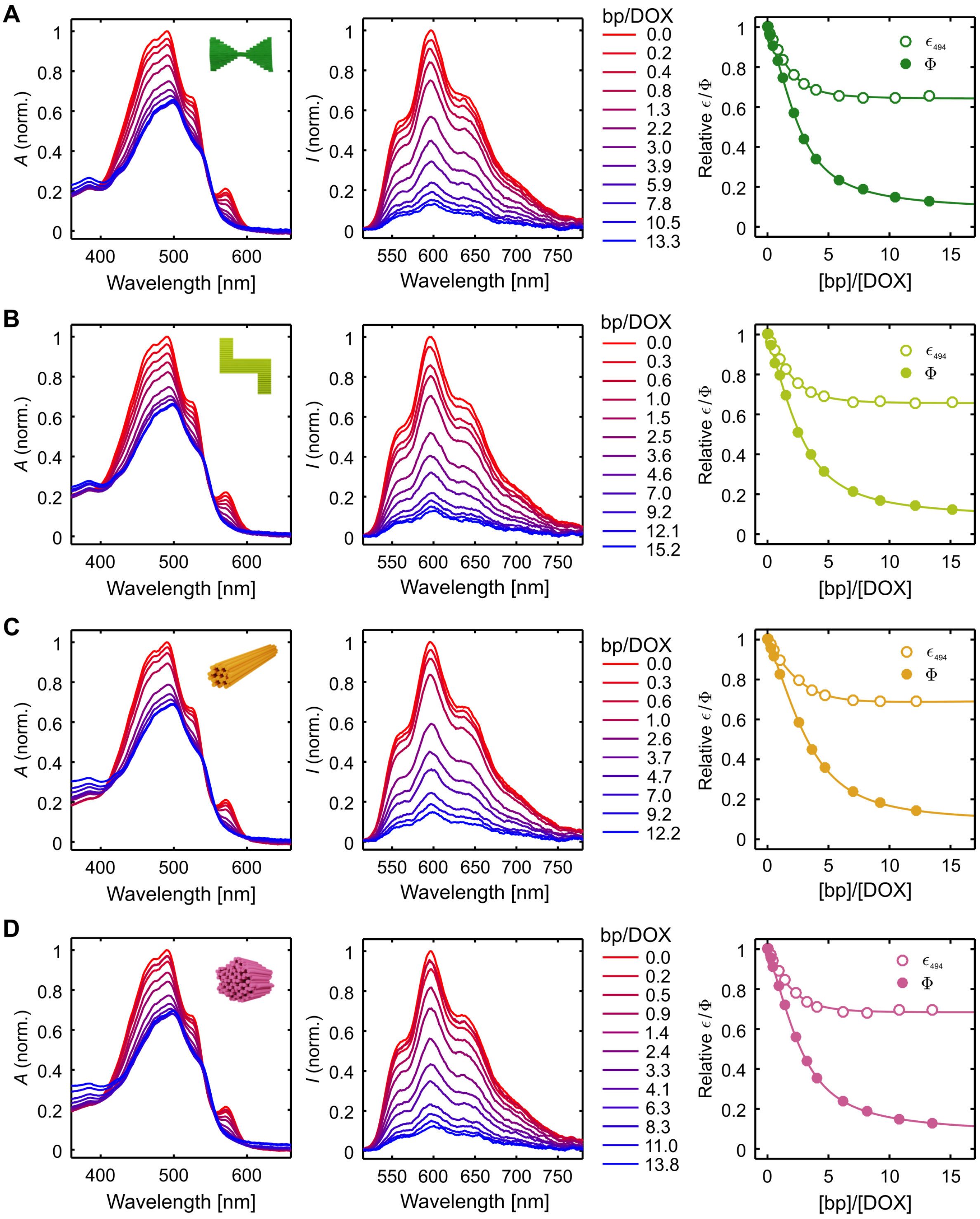

**Supplementary Figure 4.** DOX absorption spectra (left panel), fluorescence emission spectra (middle panel), and the dependence of DOX extinction coefficient at 494 nm ($\epsilon_{494}$) and fluorescence quantum yield ($\Phi$) on the amount of added DNA (right panel) for all studied DNA origami shapes. For both the absorption and emission spectra, the molar ratio of DNA base pairs and DOX (bp/DOX) is indicated in the legend. The concentration of DOX is 3 µM. The emission spectra have been collected after 494 nm excitation and corrected for the decrease of $\epsilon_{494}$. The titration isotherms on the right panel have been fitted with a 1:2 molecular binding model. The fitting parameters are listed in Supplementary Table 2. (**A**) Bowtie. (**B**) Double-L. (**C**) 24HB. (**D**) Capsule.

### Note S6: Comparison of titration isotherms ($\epsilon$ and $\Phi$) for all structures

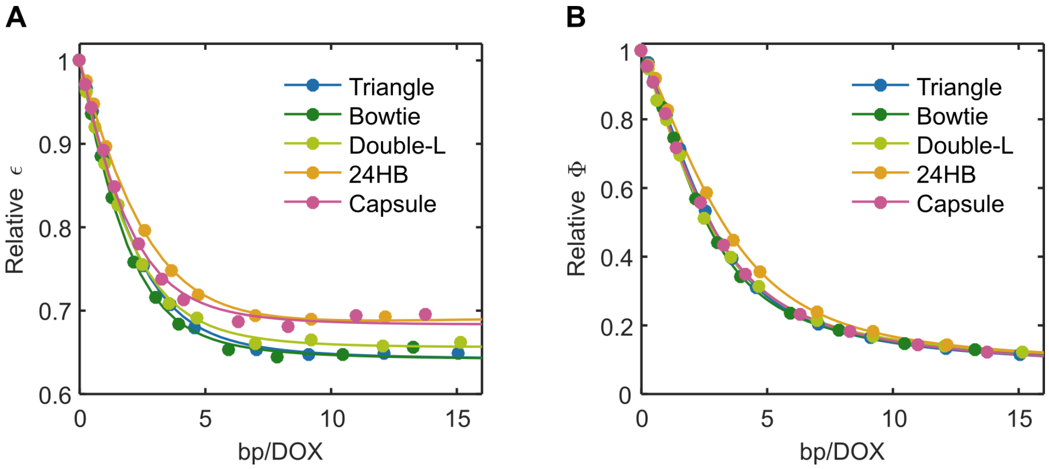

**Supplementary Figure 5.** Comparison of the effect of titrating DOX with the origami structures in terms of the relative decrease of the molar extinction coefficient at 494 nm ($\epsilon_{494}$) and fluorescence quantum yield ($\Phi$). (**A**) Relative decrease of $\epsilon_{494}$ of DOX upon addition of DNA origami. (**B**) Relative decrease of $\Phi$ measured with 494 nm excitation.

### Note S7: Density of DOX molecules in the DNA origami structures in the titration experiments

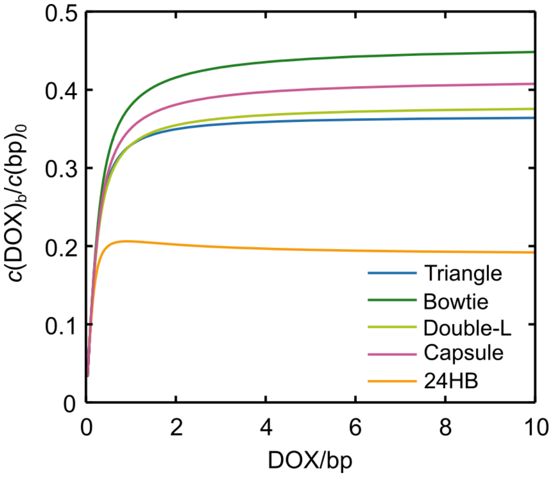

**Supplementary Figure 6.** Density of DOX molecules in the DNA origami structures in the titration experiments expressed as the number of bound DOX molecules per base pair of DNA ($c(\text{DOX})_b/c(\text{bp})_0$). The molar ratio DOX/bp on the $x$ axis refers to the total concentration of DOX and base pairs in the samples ($c(\text{DOX})_0$ and $(bp)_0$).

### Note S8: Fitting parameters $K$, $\Phi$, and $\epsilon$ for all origami shapes obtained from the 1:2 DOX-DNA binding model

**Supplementary Table 2.** Association constants ($K_{11}$, $K_{12}$), fluorescence quantum yields ($\Phi_{11}$, $\Phi_{12}$), and molar extinction coefficients at 494 nm ($\epsilon_{11}$, $\epsilon_{12}$) for the two distinct DOX-DNA complexes. The values have been obtained by fitting the titration data with the 1:2 molecular binding model as described in the Methods section. $\Phi_{11}$, $\Phi_{12}$, $\epsilon_{11}$, and $\epsilon_{12}$ are presented relative to the extinction coefficent and quantum yield of free DOX ($\epsilon_0$ and $\Phi_0$.

| Origami shape | $K_{11}$ ($\times 10^5 \mathrm{M}^{-1}$) | $K_{12}$ ($\times 10^5 \mathrm{M}^{-1}$) | $\Phi_{11}/\Phi_0$ (%) | $\Phi_{12}/\Phi_0$ (%) | $\epsilon_{11}/\epsilon_0$ (%) | $\epsilon_{12}/\epsilon_0$ (%) |
|---|---|---|---|---|---|---|
| Triangle | $1.93 \pm 0.06$ | $3.07 \pm 0.06$ | $62 \pm 3$ | $6.2 \pm 0.3$ | $62 \pm 3$ | $64 \pm 3$ |
| Bowtie | $2.81 \pm 0.12$ | $2.95 \pm 0.07$ | $64 \pm 4$ | $6.9 \pm 0.4$ | $64 \pm 4$ | $64 \pm 4$ |
| Double-L | $2.0 \pm 0.2$ | $2.15 \pm 0.14$ | $47 \pm 5$ | $7.0 \pm 0.7$ | $60 \pm 6$ | $66 \pm 7$ |
| 24HB | $0.77 \pm 0.03$ | $2.59 \pm 0.09$ | $27.8 \pm 1.2$ | $8.2 \pm 0.4$ | $42 \pm 2$ | $71 \pm 3$ |
| Capsule | $2.36 \pm 0.05$ | $2.26 \pm 0.03$ | $57 \pm 6$ | $1.3 \pm 0.2$ | $64.2 \pm 1.5$ | $69 \pm 2$ |
| **Average** | **$2.0 \pm 0.3$** | **$2.6 \pm 0.2$** | **$52 \pm 7$** | **$6.7 \pm 0.9$** | **$58 \pm 8$** | **$67 \pm 9$** |

### Note S9: AFM images of double-L loaded with DOX and DOX-loading estimation

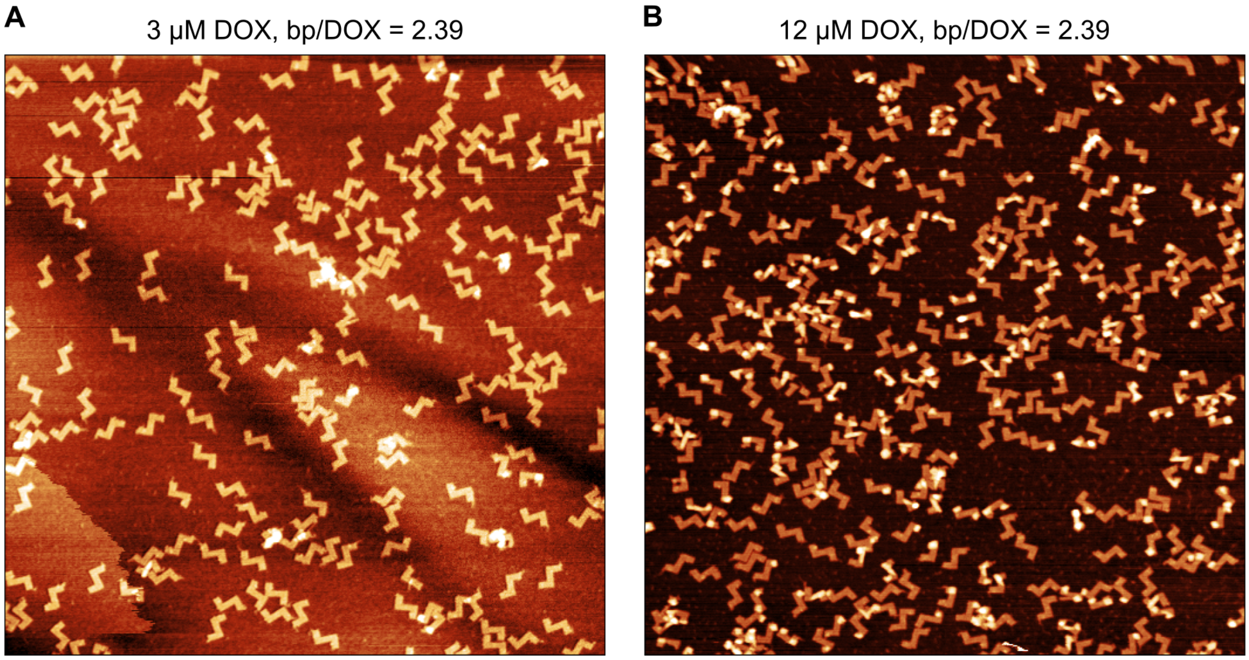

**Supplementary Figure 7.** AFM images of double-L origami loaded with DOX at two different sample concentrations while keeping the same molar ratio of DNA base pairs (bp) and DOX (bp/DOX). The edge of each figure is 3 μM in length. (**A**) DOX-origami sample prepared with a base pair concentration ($c(\text{bp})_0$) of 7.16 μM and DOX concentration ($c(\text{DOX})_0$) of 3 μM. (**B**) DOX-origami sample prepared at 4 times higher concentration of both DOX and double-L origami than Figure A ($c(\text{bp})_0$ = 28.7 μM, $c(\text{DOX})_0$ = 12 μM). While the bp/DOX ratio is identical in both cases, the higher total concentration of both DNA origami and DOX in Figure B can be seen to lead to an increased fraction of twisted DNA origami shapes, indicating higher DOX binding density through intercalation. The analysis result is in line with the theoretical values predicted by the 1:2 binding model (Supplementary Table 3.

**Supplementary Table 3.** Theoretical prediction of the density of loaded DOX (number of bound DOX molecules per base pair, $c(\text{DOX})_\text{b}/c(\text{bp})_0$) in the double-L structures shown in Supplementary Figure 7 based on the 1:2 binding model.

| | **3 μM DOX** | **12 μM DOX** |
|---|---|---|
| $c(\text{DOX})_0$ (μM) | 3.0 | 12 |
| $c(\text{bp})_0$ (μM) | 7.16 | 28.7 |
| bp/DOX | 2.39 | 2.39 |
| $c(\text{DOX})_\text{b}$ (μM) | 1.90 | 10.6 |
| $c(\text{DOX})_\text{b}/c(\text{bp})_0$ | 0.27 | 0.37 |

### Note S10: Kinetics of DOX-DNA association – comparison of different incubation times in the titration experiment

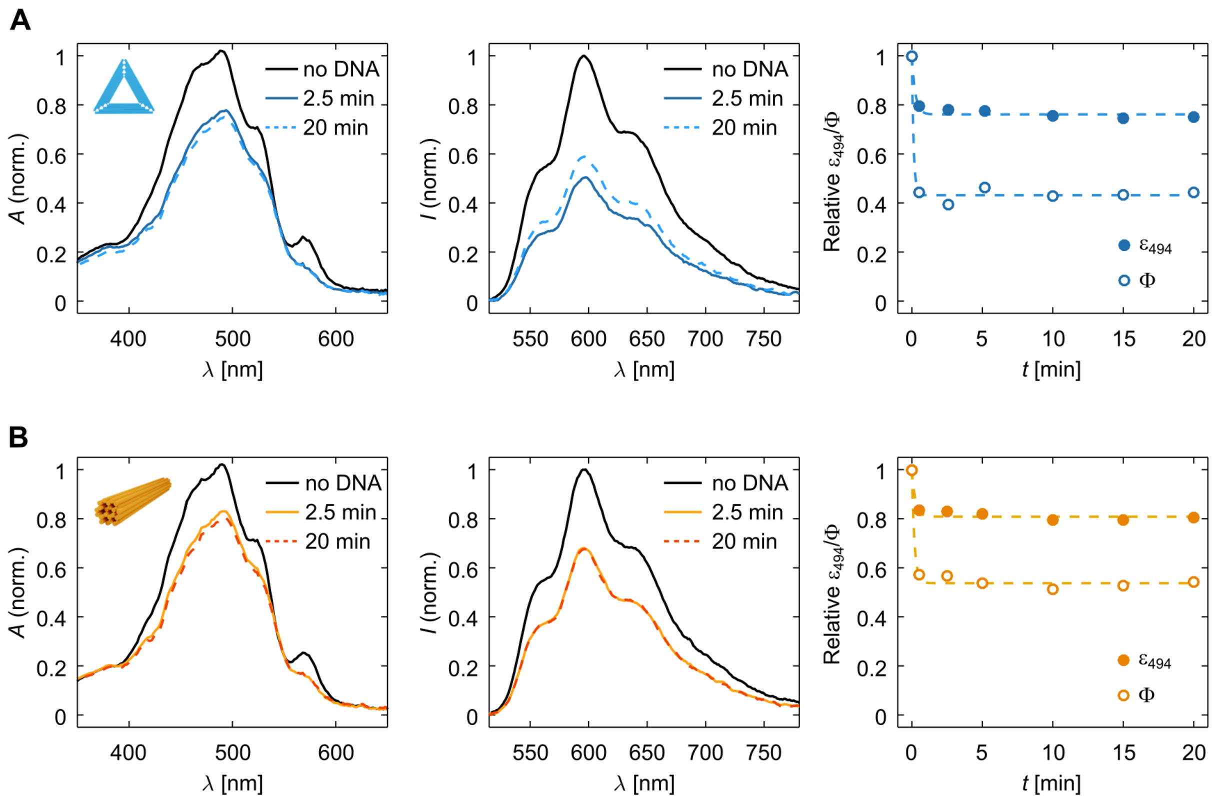

**Supplementary Figure 8.** The effect of incubation time after DNA addition on the DOX absorption and fluorescence spectra of 3 μM DOX. For confirming that the 2.5 min incubation time in the titration experiment is sufficient for reaching a binding equilibrium for all origami structures, the absorption and fluorescence spectra of 3 μM DOX were collected at different time points after addition of DONs. (**A**) DOX + triangle DON (bp/DOX = 2.16). Left panel: normalized absorption spectra of DOX in the absence of DONs, and 2.5 min and 20 min after an addition of DONs. Middle panel: Normalized fluorescence emission spectra in the absence of DONs, and 2.5 min and 20 min after an addition of DONs. The emission spectra have been collected after 494 nm excitation and corrected for the decrease of $\epsilon_{494}$. Right panel: the relative $\epsilon_{494}$ and the fluorescence quantum yield ($\Phi$) of DOX *vs.* the incubation time. (**B**) DOX + 24HB DON (bp/DOX = 1.94). The figures are presented as in Figure A.

### Note S11: Titration of DOX with ssDNA

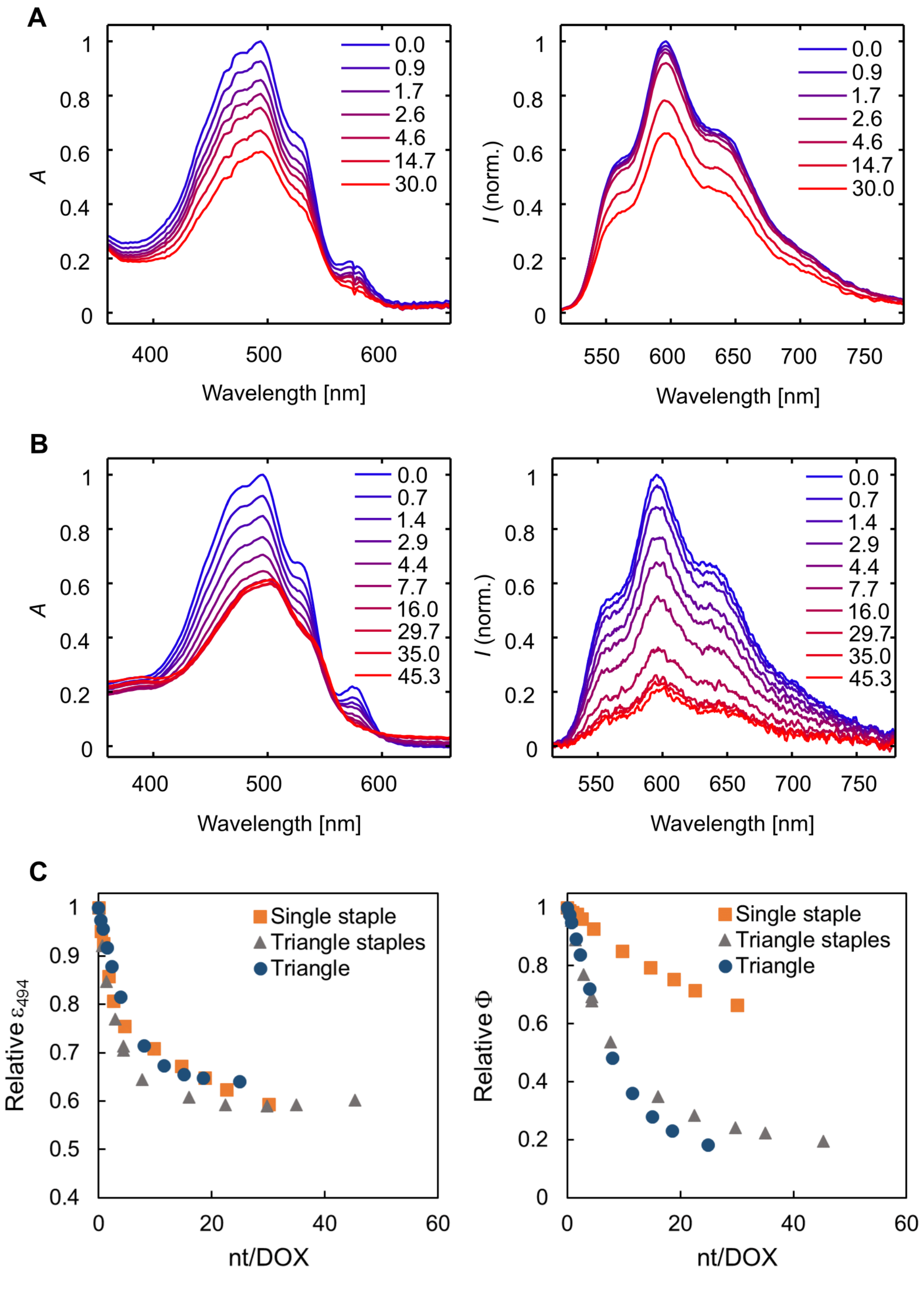

**Supplementary Figure 9.** Titration of 3 μM DOX with single- and double-stranded DNA. (**A**) Titration with a single oligonucleotide with a low probability of self-hybridization (5'–GAACAACGCTCCAACCATCGC–3'). Absorption spectrum of DOX is shown in the left panel and fluorescence emission in the right panel. The emission intensities have been collected with 494 nm excitation and corrected for the decrease of $\epsilon_{494}$ to represent the quantum yield of the DOX molecules ($\Phi$). The legends in both figures denote the molar ratio of nucleotides and DOX (nt/DOX) for each spectrum. The concentration of nucleotides ($c$(nt)$_0$ has been calculated from the measured $A_{260}$ value and a molar extinction coefficient for 100% single-stranded DNA, $\epsilon_{260}$ = 10,000 M$^{-1}$cm$^{-1}$. (**B**) Titration with a mixture of the 232 staple oligonucleotides used for folding the triangle origami, presented as Figure A. While the mixture of staple strands can be expected to contain a high number of hybridized dsDNA regions, the exact fraction of ssDNA and dsDNA nucleotides in the sample is unknown. The nt/DOX ratios presented in the legends have been calculated from the measured $A_{260}$ value and a molar extinction coefficient for 100% single-stranded DNA, $\epsilon_{260}$ = 10,000 M$^{-1}$cm$^{-1}$. (**C**) The relative decrease of $\epsilon_{494}$ and $\Phi$ compared upon titration with a single oligonucleotide (Figure A), the triangle staple oligonucleotide mixture (Figure B), and the folded triangle origami (Figure 3B in the main text).

### Note S12: Scattering intensity of the origami

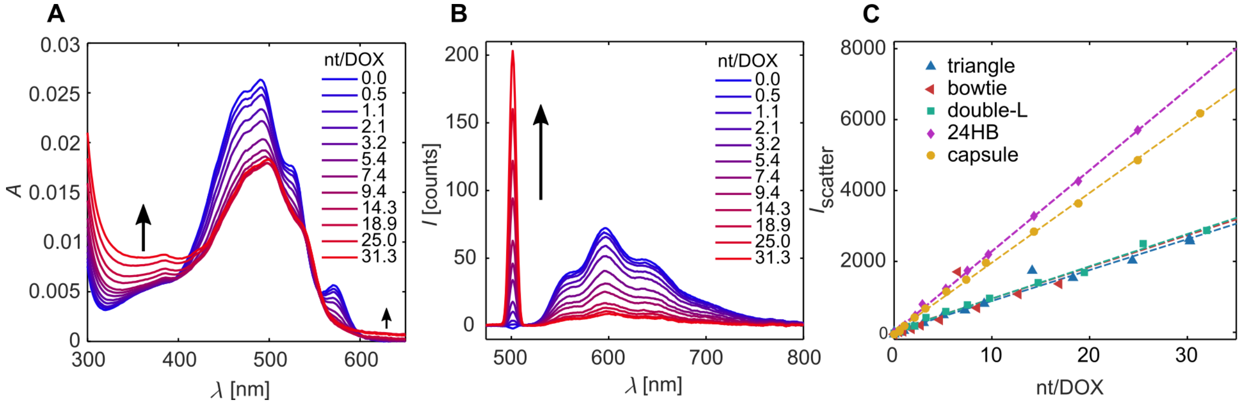

**Supplementary Figure 10.** Light scattering caused by the DOX-loaded DNA origami. (**A**) Absorption spectrum of a sample containing DOX and an increasing concentration of the capsule origami. Absorption baseline shift upwards during the titration is indicated with the black arrows and shows the increased light scattering in the sample. The amount of DNA origami is described as the total concentration of nucleotides in the sample, and indicated in the legend as the molar ratio of nucleotides vs. DOX (nt/DOX) ($c$(DOX) = 3 μM). (**B**) Scattered light detected as an increasing intensity of excitation light (494 nm) in the fluorescence emission measurement (90° detection relative to the excitation beam). The spectra are collected for 3 μM DOX upon titration with the capsule origami. (**C**) Intensity of scattered light during titration of DOX with different origami shapes (increasing nt/DOX ratio) shows the stronger light scattering detected with the 3D structures. The scattering intensities have been obtained from the emission spectra by integrating the excitation light peak.

### Note S13: Purification of free DOX from DOX-DON samples after loading – comparison of spin-filtration and PEG precipitation

In the experiments presented in the main article, we studied the use of centrifugation as a method for removing free DOX from DOX-loaded DONs after the system has reached an equilibrium and the loading has been completed. As the purification of free DOX from DOX-loaded DONs is an important sample preparation step for downstream applications, we studied it further by performing additional purification experiments. Here, we evaluated the efficiency of other purification methods for free DOX removal: spin-filtration and PEG precipitation.

As an initial experiment for optimizing the spin-filtration process, we studied the suitability of spin-filters with different molecular weight cutoff membranes for DOX removal. The experiment was performed by centrifuging samples of partly aggregated DOX (2 mM DOX solution prepared in FOB pH 8.0 and incubated for 24 h at RT) through 10 kDa, 30 kDa, and 100 kDa spin-filters (Figure S11a). For both 10 kDa and 30 kDa filters, the DOX concentration in the filter increased during centrifugation [$c$(concentrate) > $c_0$], which indicates that some of the DOX particles were too large to pass through the membrane (Figure S11b). Only the 100 kDa filters had sufficiently large pore size to allow the DOX molecules and aggregates to freely pass through the membrane during centrifugation, indicated by a similar DOX concentration in the unpurified sample and in the concentrate [$c_0 \approx c$(concentrate)]. In conclusion, this shows both that DOX aggregates formed during long incubation times can be relatively large in size (too large to pass through the 10 kDa and 30 kDa filter membranes), but that they can still be removed with 100 kDa molecular weight cut-off spin-filtration.

However, the DOX concentration in the filtrate [$c$(filtrate)] decreased relative to $c_0$ for all tested filters. The filter membranes also turned visibly red during the spin-filtration process. Together these observations show that part of DOX is lost in the spin-filtration process due to binding to the filter membrane. For 2 mM DOX and 100 kDa membrane, $\sim$20% of the DOX molecules were bound to the membrane during centrifugation. The extent of DOX binding to the 100 kDa filter membrane was studied also for smaller DOX concentrations and both DOX-DON and DOX-only samples. As shown in Figure S11c, the percentage of DOX binding to the filter membrane can be considerable especially at low DOX concentrations (> 90% of DOX bound to the membrane at 3 µM and 6 µM DOX concentration) – regardless of the DON concentration in the sample.

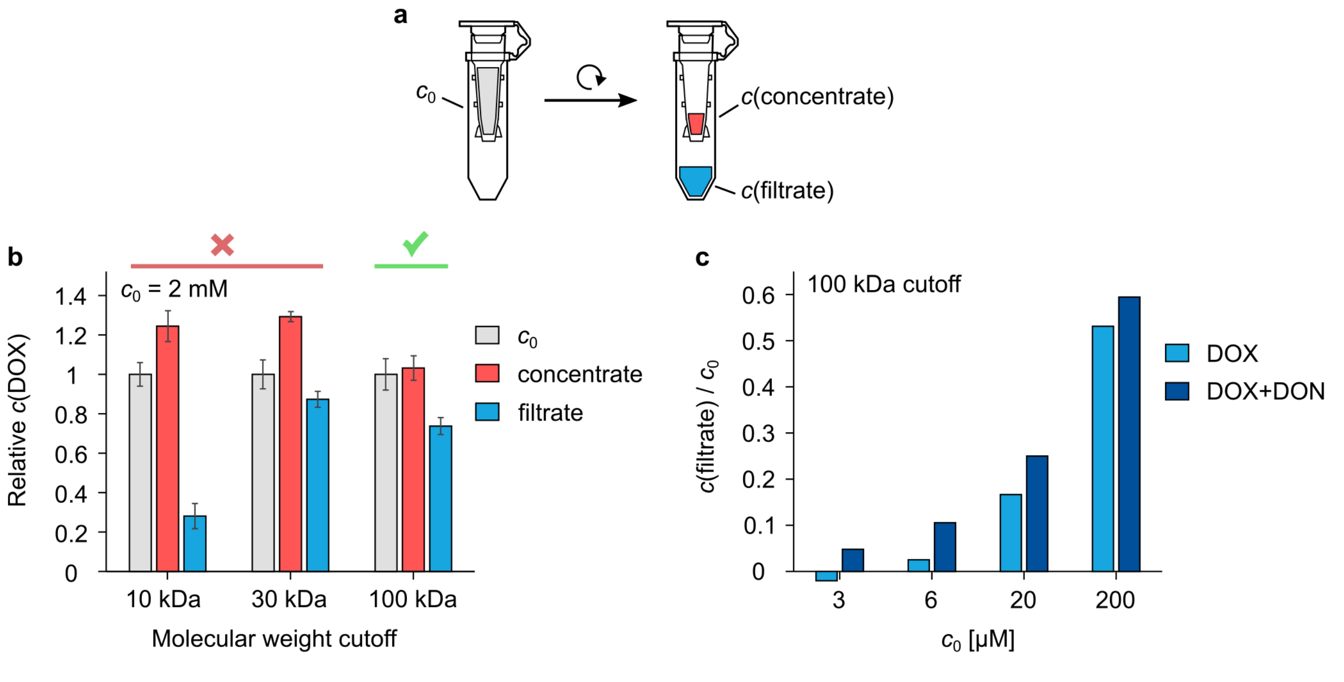

**Supplementary Figure 11.** Experiments for estimating the DOX aggregate size and the suitability of spin-filtration as a purification method for free DOX. (**a**) Schematic of the spin-filtration experiments. $c_0$ refers to DOX concentration in the sample before spin-filtration. After centrifugation, the concentration of DOX is measured from the sample remaining in the filter [$c$(concentrate)] and/or from the flow-through [$c$(filtrate)]. (**b**) Spin-filtration of 2 mM DOX solution in FOB pH 8.0 through different molecular weight cutoff filters after 24 h incubation at RT to promote aggregate formation. (**c**) $c$(filtrate) relative to $c_0$ in DOX and DOX-DON samples after spin-filtration through a 100 kDa membrane, showing the fraction of DOX bound to the filter membrane during the spin-filtration procedure. DOX samples contain the indicated $c_0$ of DOX in Tris/$Mg^{2+}$ pH 7.4. The DOX-DON samples contain additionally 2 nM of the triangle DON. All samples have been incubated for 1 h at RT before spin-filtration to load the DONs with DOX. The experiment in Figure b was repeated three times and the results are expressed as the mean ± standard deviation. The experiment in Figure c was performed once.

We then prepared DOX-loaded triangle DONs at different DOX loading concentrations (2 nM triangle DON incubated for 1 h with either 3, 6, or 20 μM DOX in Tris/$Mg^{2+}$ pH 7.4) and studied the purification outcome of both 100 kDa spin-filtration and PEG precipitation. The purification protocols have been described in the Supplementary Methods.

Both spin-filtration and PEG precipitation are highly effective for purifying free DOX from the solution (> 99% of free DOX removed with both methods). but spin-filtration leads to notably higher sample loss during purification with all DOX loading concentrations (Figure S12a). Interestingly, increasing the amount of DOX weakens the outcome of both purification methods: for spin-filtration, a clear decrease of sample recovery is observed with increasing DOX concentration. PEG precipitation leads to high sample recovery with all DOX concentrations, but the formed precipitates were observed to become increasingly difficult to resuspend at higher DOX concentrations. While 3 μM and 6 μM pellets were dissolved after overnight incubation at RT, the 20 μM DOX concentration required additional incubation under shaking at +30 °C to dissolve. As this could cause some sample damage, the results for PEG-purification at 20 μM DOX concentration may not be fully comparable to the other loading concentrations, indicated with a different color in the presented figures.

We then studied the composition of the purified samples by determining their DNA and DOX concentrations with UV-Vis analysis. Because free DOX has been removed from the samples, the analysis can be used to determine the DOX loading density (DOX/bp) – number of bound DOX molecules per a DNA base pair – for each DOX loading concentration. This analysis thus also complements the results of DOX loading yield obtained by studying the DOX-DON binding equilibrium with titration experiments as well as the loading yield estimation in the DNase I experiment. As seen in Figure S12b, the DOX/bp ratio increases with the DOX loading concentration, but in a non-linear fashion. This is an expected result based on the DOX-DNA binding equilibrium and in line with the titration experiments, where we estimated the DONs to be saturated with DOX at DOX/bp = 0.36 ± 0.10. The solid line in Figure S12b presents the equation

$$\mathrm{DOX/bp} := \frac{c(\mathrm{DOX})_\mathrm{b}}{c(\mathrm{bp})_0} = \frac{c(\mathrm{DOX})_0}{c(\mathrm{bp})_0} \times s \times f_b \tag{1}$$

Where $f_\mathrm{b}$ for each $c(\mathrm{bp})_0$ is obtained from Equations (3) and (4) in the main article using $K_{11}$ = (2.0 ± 0.3) $\times$ $10^5$ $\mathrm{M}^{-1}$ and $K_{12}$ = (2.6 ± 0.2) $\times$ $10^5$ $\mathrm{M}^{-1}$. As a feature of the applied binding model, the DOX/bp ratio approaches 1 at high $c(\mathrm{DOX})_0$. A factor $s$ (the DOX/bp saturation value) was thus included in the Equation 1 for correcting the model for a saturation value < 1 by finding a value of $s$ that leads to the best fit to the data with non-linear least-squares fitting. The best fit was obtained with $s$ = $(\mathrm{DOX/bp})_\mathrm{max}$ = 0.45 ± 0.04 DOX/bp.

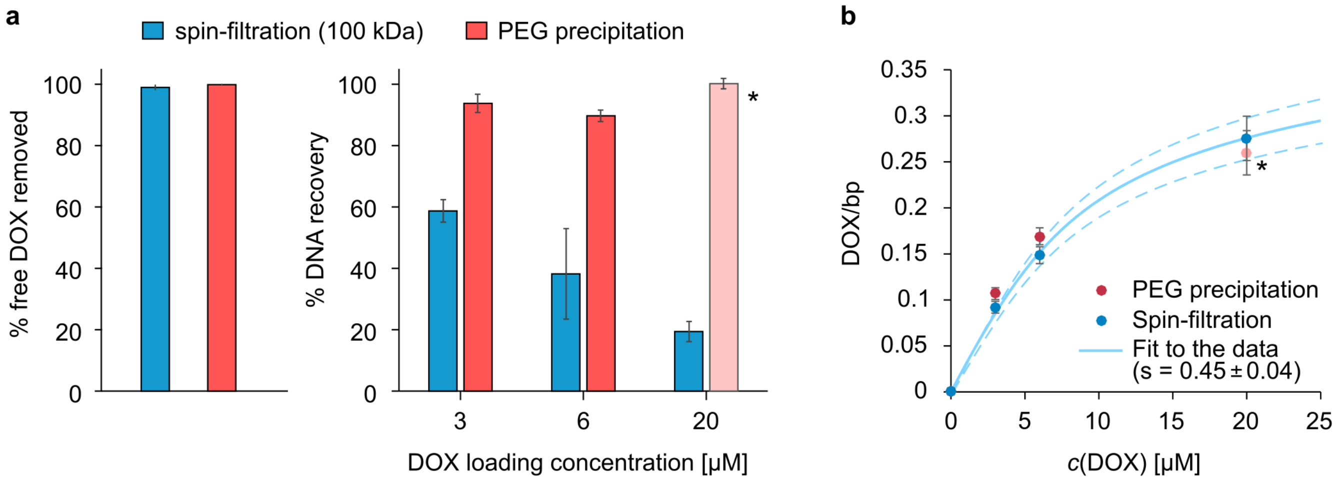

**Supplementary Figure 12.** Comparison of spin-filtration (100 kDa) and PEG precipitation as purification methods for removing free DOX from DOX-loaded triangle DONs. (**a**) DOX purification (free DOX removal) efficiency and sample loss during the purification. *PEG purification result of the 20 μM sample is shown, although we note that PEG precipitation at this DOX concentration is not recommended as the resulting pellet required heavy processing to dissolve. (**b**) DOX-DON sample composition after purification. The number of bound DOX molecules per DNA base pair (DOX/bp) was determined from purified DOX-DON samples prepared at 3–20 μM DOX loading concentration. The best fit of the thermodynamic binding model to the data is obtained with a saturation at 0.45 ± 0.04 DOX/bp.

#### Note S14: DOX release from DONs through diffusion after purification

After confirming that spin-filtration efficiently removes free DOX from DOX-loaded DON samples (Supplementary Note S13), we studied the kinetics of the diffusive release of DOX from the DONs. This takes place when the low-DOX environment of the solution favors the dissociation of DOX (Supplementary Figure 13a). Through a series of experiments described below, we conclude that the release of DOX from DONs takes place rapidly after the binding equilibrium is disturbed by removing the unbound DOX molecules. We did not explicitly study the kinetics of the release, but could define that freshly spin-filtered, DOX-loaded bowtie DON samples had already reached a new binding equilibrium in less than 10 minutes after the spin-filtration procedure.

Supplementary Figure 13b shows the change of the base pair concentration [$c(\mathrm{bp})_0$] and DOX concentration [$c(\mathrm{DOX})_0$] when DOX-loaded bowtie samples (2 nM DONs + 3 or 6 μM DOX) were either purified with 100 kDa spin-filtration or diluted to the same $c(\mathrm{bp})_0$ as in the purified sample. The purification of unbound DOX was expected to lead to a considerably reduced observed quantum yield of DOX ($\Phi_{\mathrm{obs}}$) when the remaining DOX molecules are bound to DONs. Still, only a slight decrease of $\Phi_{\mathrm{obs}}$ was observed in the purified sample, indicating a considerable number of free, fluorescent DOX molecules (Supplementary Figure 13c). As the efficiency of the purification of free DOX molecules was confirmed to be ∼100% (Supplementary Figure 12a), it can be assumed that the free DOX molecules were released from the DONs after the purification. A dilution of the unpurified samples to the same $c(\mathrm{bp})_0$ as in the purified samples also led to DOX release, observed as a slight increase of the $\Phi_{\mathrm{obs}}$.

In a further analysis, the detected values of $\Phi_{\mathrm{obs}}$ were compared to the expected quantum yields after equilibration ($\Phi_{\mathrm{eq}}$) (Supplementary Figure 13c). The $\Phi_{\mathrm{eq}}$ values were calculated from the Equation 2 for each $c(\mathrm{DOX})_0$ and $c(\mathrm{bp})_0$ according to the applied binding model and the values of $K_{11}$, $K_{12}$, $\Phi_{11}$, and $\Phi_{12}$ for the bowtie DON (Supplementary Table 2). A close agreement between the measured and calculated values was observed. This indicates that the DOX release from the purified samples had progressed to a thermodynamic equilibrium before the measurement (in less than 10 minutes after spin-filtration). The loading density for the samples in terms of DOX/bp is obtained by dividing the bound DOX concentration $c(\mathrm{DOX})_{\mathrm{b}}$ obtained from the binding model (Equation 4) by the $c(\mathrm{bp})_0$. This shows that the purification and subsequent DOX release decreased the DOX loading density of DONs by 70% for 3 μM and by 66% for the 6 μM loading concentration (Supplementary Figure 13d). No further $\Phi_{\mathrm{obs}}$ increase was detected when the spin-filtered samples were incubated at RT for 45 hours (Supplementary Figure 13e). This indicates that the equilibrium state had been reached before the first measurement. When the diluted and the purified DOX-bowtie samples were subjected to DNase I digestion, the reduced total DOX concentration in the purified samples led to increased digestion and DOX release rates (Supplementary Figure 14).

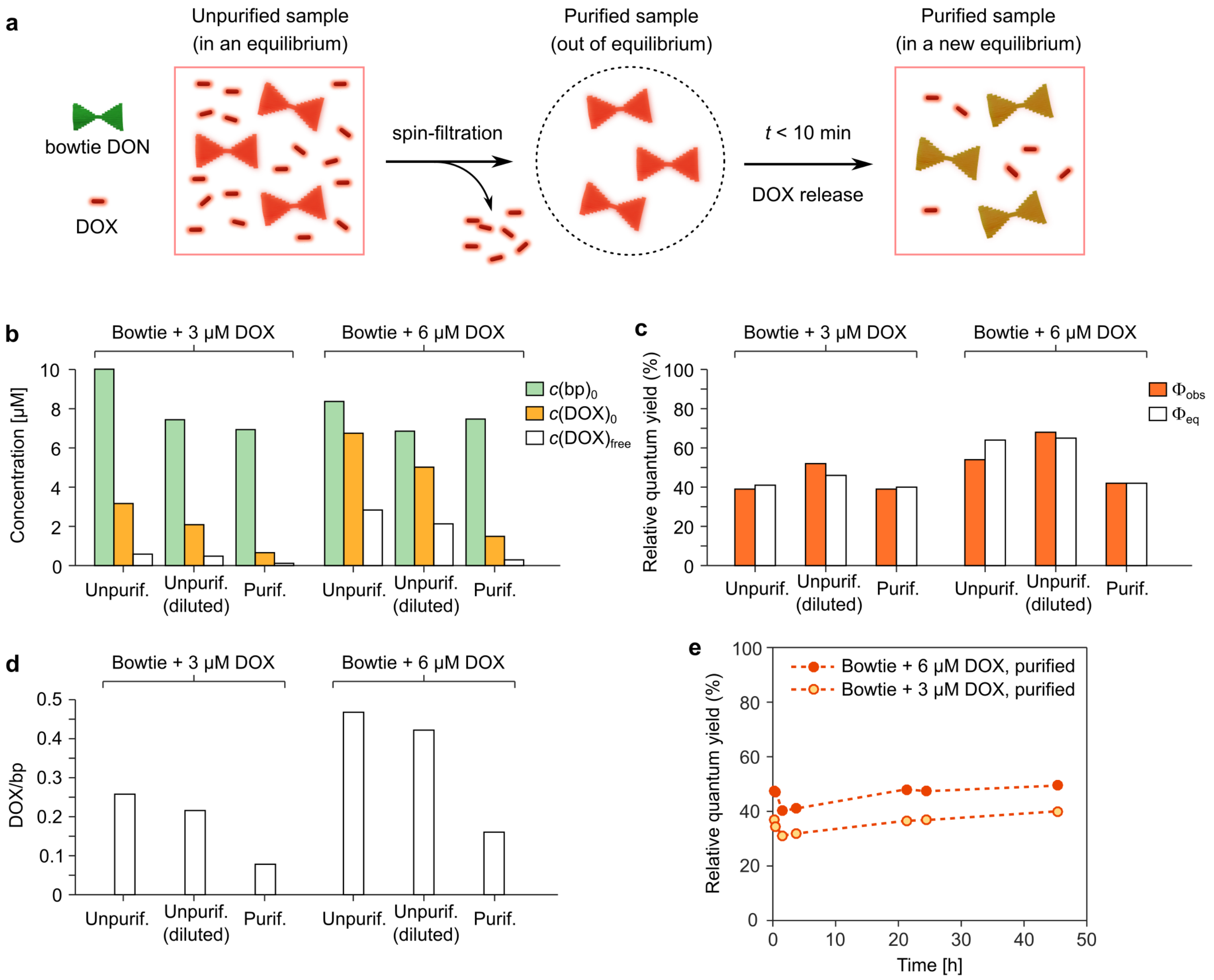

**Supplementary Figure 13.** Evidence of a fast equilibration and DOX release after free DOX removal from DOX-loaded bowtie DONs. (**a**) Schematic illustration of the sample composition after the spin-filtration and the subsequent equilibration, during which a part of the bound DOX molecules is released from the DONs. (**b**) The measured base pair concentrations [$c$(bp)$_0$] and total DOX concentrations [$c$(DOX)$_0$] in the studied DOX-bowtie samples: unpurified samples, unpurified but diluted samples, and spin-filtrated samples. The calculated equilibrium state free DOX concentrations [$c$(DOX)$_{\mathrm{free}}$] for each sample are also shown. In subfigures b, c, and d, colored bars represent experimentally determined values, while empty bars are values calculated according to the applied binding model (Equations 2–4 in the main text) from the experimental values. (**c**) The measured DOX quantum yield ($\Phi_{\mathrm{obs}}$) in freshly spin-filtered DOX-bowtie samples (< 1 h) compared to the calculated equilibrium state quantum yields ($\Phi_{\mathrm{eq}}$). The close correspondence between the measured and calculated values suggests that all three sample types have reached an equilibrium before the measurement. (**d**) The calculated DOX loading densities in terms of bound DOX molecules per DNA base pair (DOX/bp). (**e**) The detected $\Phi_{\mathrm{obs}}$ of spin-filtered samples during 45-hour incubation. Subfigures b, c, and d show representative data obtained for a set of samples in a single experiment. Qualitatively reproducible results were obtained when the experiment was repeated.

### Note S15: The DNase I digestion and DOX release profiles of bowtie DON samples before and after spin-filtration

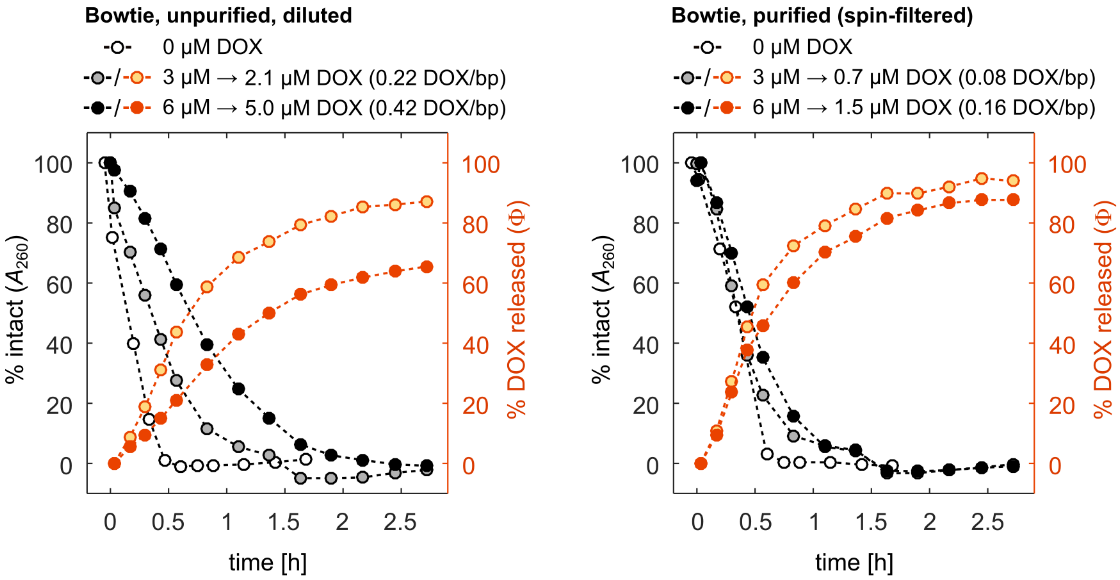

**Supplementary Figure 14.** The effect of spin-filtration on the the DNase I digestion profiles (white, gray, and black circles) and DOX release (yellow and orange circles) of bowtie DON samples. On the left, DOX-loaded bowtie samples prepared at either 3 or 6 μM DOX loading concentration were diluted before the digestion to a match the base pair concentration of the purified samples. On the right, the samples were purified with spin-filtration to remove the originally unbound DOX before the digestion. As described under Supplementary Note S14, all samples were in a thermodynamic equilibrium before starting the digestion and contain both bound and free DOX molecules as presented in Supplementary Figure 13b. The presented data was obtained for a set of samples in a single experiment. Qualitatively reproducible results were obtained when the experiment was repeated.

### Note S16: Details of DOX-DON samples in the DNase I digestion experiments

**Supplementary Table 4.** Composition of the DNA origami-DOX samples in the DNase I digestion experiments before addition of DNase I. The DOX quantum yields relative to free DOX references ($\Phi/\Phi_{\mathrm{ref}}$) and the base pair/DOX molar ratios (bp/DOX) are based on the measured DNA absorbance and DOX fluorescence intensities. The concentrations of free DOX ($c$(DOX)$_{\mathrm{ub}}$), bound DOX ($c$(DOX)$_{\mathrm{b}}$), and bound DOX molecules bound per base pair of DNA ($c$(DOX)$_{\mathrm{b}}/c$(bp)$_0$) have been calculated according to Equation 6 and the parameters in Supplementary Table 2. All values are presented as the mean $\pm$ standard error of three individual measurements.

**3 μM DOX**

| | $\Phi/\Phi_{\mathrm{ref}}$ | bp/DOX | $c$(DOX)$_{\mathrm{ub}}$ (μM) | $c$(DOX)$_{\mathrm{b}}$ (μM) | $c$(DOX)$_{\mathrm{b}}/c$(bp)$_0$ |
|---|---|---|---|---|---|
| Triangle | $0.37 \pm 0.06$ | $4.0 \pm 0.3$ | $0.68 \pm 0.11$ | $2.3 \pm 0.2$ | $0.196 \pm 0.015$ |
| Bowtie | $0.44 \pm 0.13$ | $3.7 \pm 0.2$ | $0.9 \pm 0.4$ | $2.1 \pm 0.4$ | $0.19 \pm 0.04$ |
| Double-L | $0.43 \pm 0.13$ | $3.6 \pm 0.2$ | $1.0 \pm 0.4$ | $2.0 \pm 0.4$ | $0.19 \pm 0.04$ |
| Capsule | $0.48 \pm 0.12$ | $4.9 \pm 0.5$ | $0.9 \pm 0.5$ | $2.1 \pm 0.5$ | $0.14 \pm 0.02$ |
| 24HB | $0.33 \pm 0.05$ | $5.3 \pm 0.6$ | $0.9 \pm 0.3$ | $2.1 \pm 0.3$ | $0.14 \pm 0.01$ |

**6 μM DOX**

| | $\Phi/\Phi_{\mathrm{ref}}$ | bp/DOX | $c$(DOX)$_{\mathrm{ub}}$ (μM) | $c$(DOX)$_{\mathrm{b}}$ (μM) | $c$(DOX)$_{\mathrm{b}}/c$(bp)$_0$ |
|---|---|---|---|---|---|
| Triangle | $0.55 \pm 0.05$ | $1.98 \pm 0.15$ | $2.4 \pm 0.2$ | $3.6 \pm 0.2$ | $0.31 \pm 0.04$ |
| Bowtie | $0.62 \pm 0.13$ | $1.86 \pm 0.10$ | $3.0 \pm 0.8$ | $3.0 \pm 0.8$ | $0.28 \pm 0.09$ |
| Double-L | $0.60 \pm 0.13$ | $1.80 \pm 0.10$ | $3.2 \pm 0.6$ | $2.8 \pm 0.6$ | $0.27 \pm 0.07$ |
| Capsule | $0.63 \pm 0.13$ | $2.4 \pm 0.3$ | $2.7 \pm 1.0$ | $3.3 \pm 1.0$ | $0.21 \pm 0.05$ |
| 24HB | $0.61 \pm 0.12$ | $2.6 \pm 0.3$ | $2.7 \pm 0.9$ | $3.3 \pm 0.9$ | $0.20 \pm 0.04$ |

### Note S17: Comparison of the spectroscopic results and an agarose gel electrophoresis (AGE) analysis of the DNase I digestion

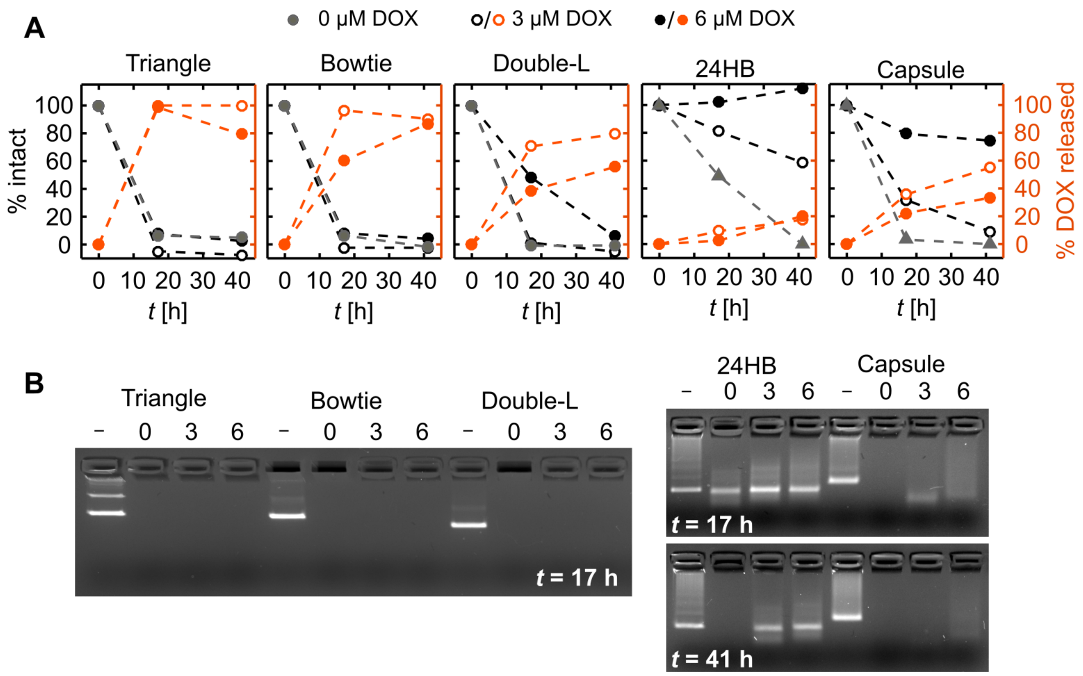

**Supplementary Figure 15.** Parallel analysis of the DNase I digestion of DONs with both spectroscopic means and with an agarose gel electrophoresis (AGE). 28.2 U/mL DNase I has been added in all samples at *t* = 0. (**A**) Digestion and DOX release profiles based on the $A_{260}$ readings (% intact) and the recovery of DOX quantum yield (% DOX released). The structural integrity of the DONs is shown with the gray markers for the samples without DOX, and with the empty and filled black markers for the samples containing DOX at either 3 or 6 $\mu$M concentration. For the samples with 3 and 6 $\mu$M DOX, the empty and filled orange markers depict the fraction of released DOX molecules relative to the initial concentration of bound DOX molecules. (**B**) AGE result of the 2D DONs after 17 h digestion and 3D DONs after 17 h and 41 h digestion. The first lane for each sample (–) contains intact DONs in the absence of DNase I at 0 μM DOX concentration; the lanes marked with 0, 3, and 6 contain the digested DONs with the indicated concentration of DOX (in $\mu$M).

### Note S18: 24HB design and staple sequences

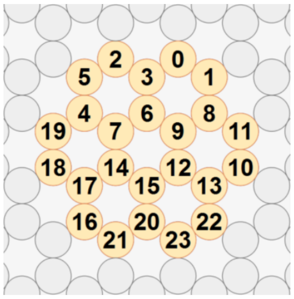

**Supplementary Figure 16.** Cross-section of the 24 helix bundle (24HB) design and the helix numbers.

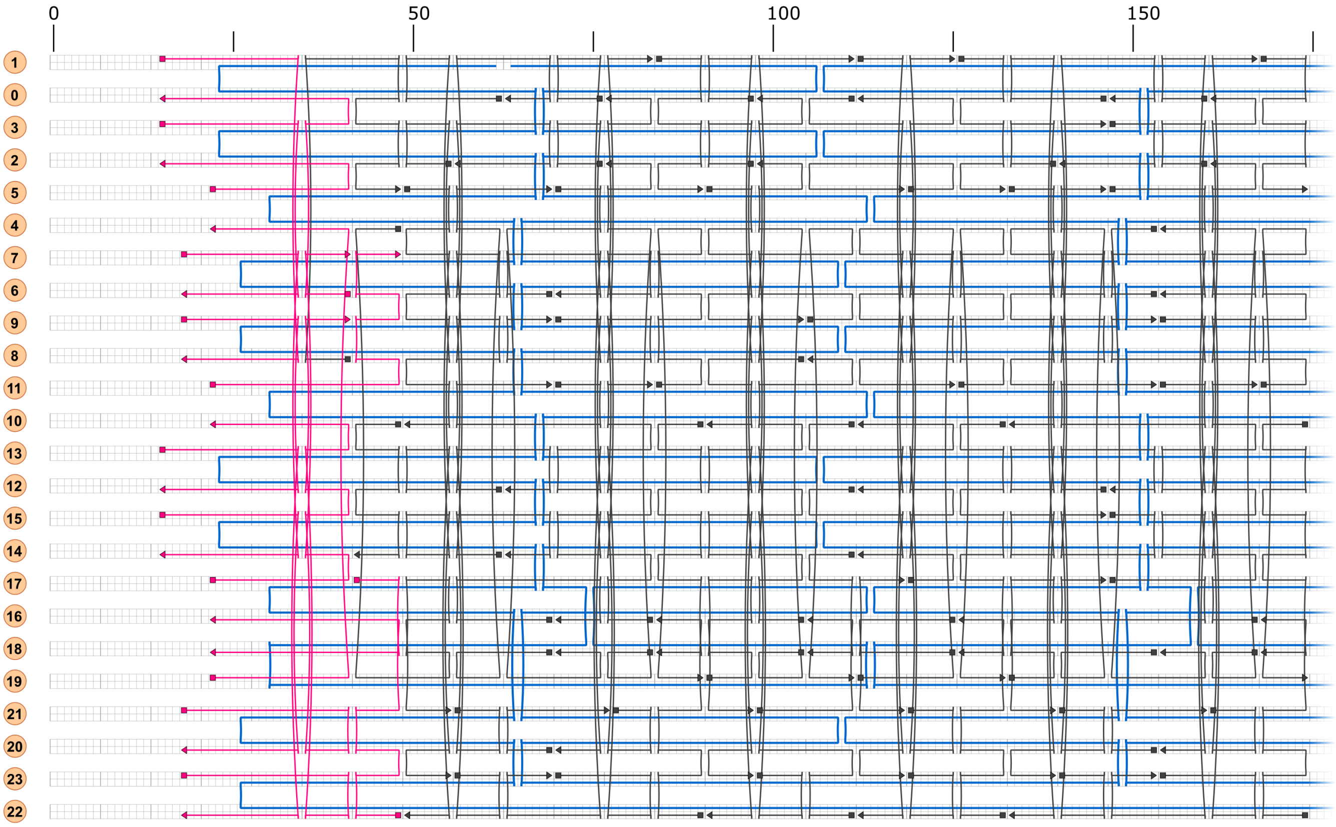

**Supplementary Figure 17.** Part 1 (grid positions 15–174) of the 24HB CaDNAno blueprint. The helix numbers are shown on the left side of the figure, and the numbers on top indicate the grid position. The 7,560-nt scaffold is shown in blue, fully complementary scaffold strands in dark gray, and scaffold strands containing 8× poly-T overhang sequences in pink.

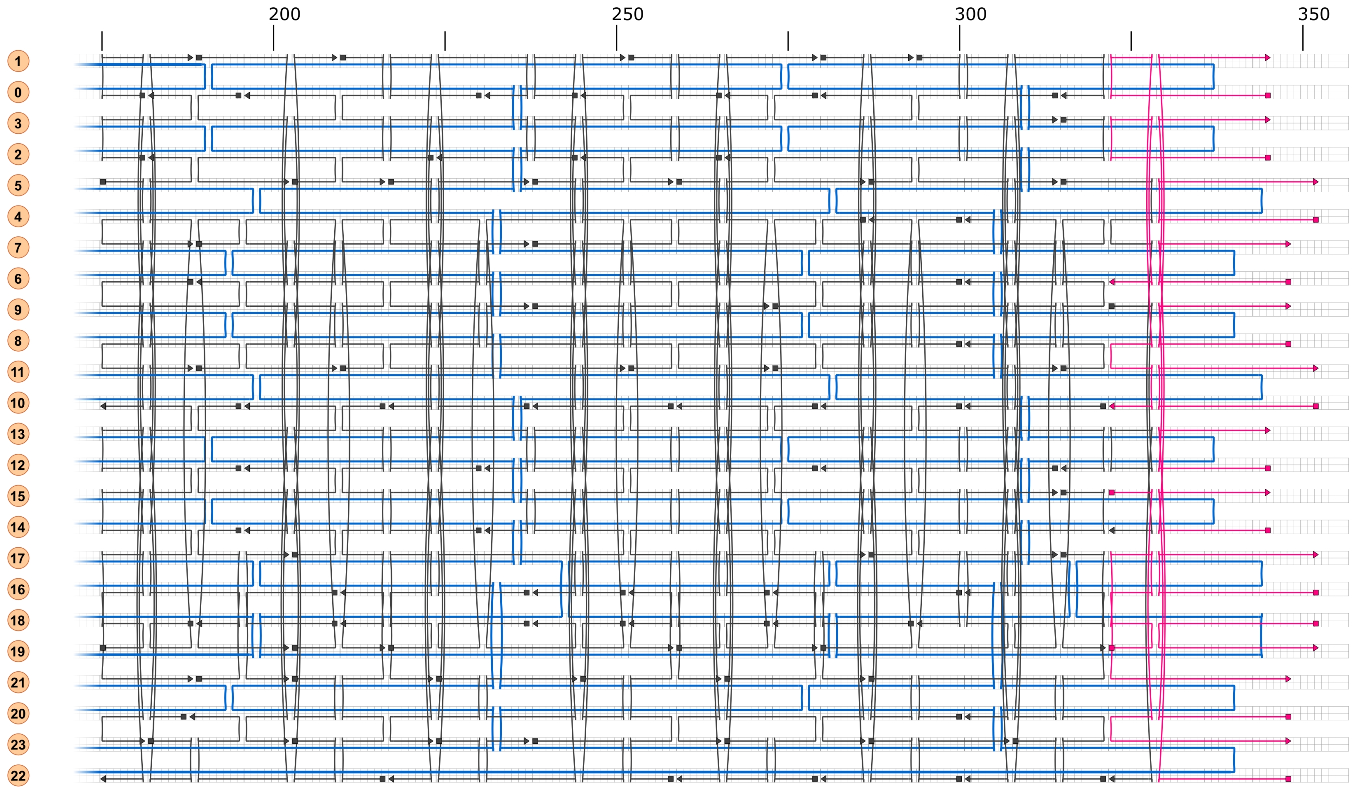

**Supplementary Figure 18.** Part 2 (grid positions 175–352) of the 24HB CaDNAno blueprint.

**Supplementary Table 5.** Staple strand sequences for the 24HB design. Nucleotide bases complementary to the scaffold strand are written in capital letters. The 8× poly-T overhang sequences in staples #174–202 have been indicated by lowercase letters. Start pos. and end pos. indicate the positions of the staple strand termini in the caDNAno design (helix number[grid position]).

| # | Start pos. | End pos. | Sequence |
|---|---|---|---|
| 1 | 3[147] | 2[140] | TTGAAATTCAAATCCAATCGC |
| 2 | 6[153] | 20[154] | CTCATTTATCTAATTTACGAGAGAGAGAATTTATCAAAGACA |
| 3 | 22[279] | 9[272] | GACAGTAGTTGGGAGAGGCTTGAGATGGGCAAAAGGGAAGTTAGCTTGA |
| 4 | 21[78] | 1[83] | GAAGGTAGATAATCAGTCACGGATTATATCCTGAGCAAATTTACGTGA |
| 5 | 1[210] | 5[216] | GTACCGCTAAGTATTTTTGCTTTAAGAGGCCCCCT |
| 6 | 5[238] | 18[238] | TTTTGTCGAGAATAGAAAGTACCGGAACGAGGCGCATGAAAC |
| 7 | 10[48] | 14[42] | TATCCGCTCGAATTGTGCTTGGACCTCCTTGAATC |
| 8 | 22[258] | 16[252] | ATCGCACGCTCATTTCAACTTGAAACACACGTAAC |
| 9 | 5[70] | 18[70] | GGCGCTAGCGTACTATGGTCAAATACCGAACGAACGCACGTG |
| 10 | 10[195] | 20[188] | GAAGCCTTTCCAGAGCCACCAGCCATTCGCCATTCGGTTTACAAATACAT |
| 11 | 5[287] | 5[286] | CGGAAGCTTAATTGCTCCTTTTGATAATTCTGTATGGCAGAC |
| 12 | 7[238] | 21[244] | ACTAAAGCGATTATACGAAGGTGTGAATTACCTAGGAATAAGGCTTGCC |
| 13 | 20[153] | 21[160] | CCACGACGCTTAAGACTCCTT |
| 14 | 17[287] | 0[280] | CCAAAAGATAACCCAAGACTTTCAAAAATGCGGATTAGCTCAACATGTT |
| 15 | 21[140] | 15[146] | GATGATGATTATCATACCTTT |
| 16 | 19[217] | 21[223] | TCCATGTATTTGTAGTCACCAAGCAAAACTTGAGC |
| 17 | 21[189] | 7[188] | AATACAGACCGAGGAAACGCACAAAGTTCGTCAAATCGGCTG |
| 18 | 0[195] | 12[196] | GTATCAGAGAGGATTAGGATTCGTCATACAAACAAAAATCAC |
| 19 | 2[265] | 22[259] | ACAGCCCCTACAACAGGCTCCCTTAAACTCCATTACTAAAGACAGGAAG |
| 20 | 0[181] | 10[175] | TTGAGAAGCCAACGCAAGCAAAACCTCCAGATTAG |
| 21 | 2[181] | 22[175] | AGTACCGTCGAGCCAAGAACGTCATCGTAATTTGCGCTAACGGCAACTG |
| 22 | 16[83] | 0[77] | AGTTGGCCAAATGATTCTGACCACGTATAAGGGATCCCGATTCTAAATC |
| 23 | 18[167] | 2[161] | CCCTTTTAAGAATTAGAAAAAACAATAAGTAAAGT |
| 24 | 1[126] | 5[132] | AAAGCCTTAAGGCGTTCTGACAGAACGCATATAAC |
| 25 | 4[300] | 16[301] | TTCGAGCTTGATAATCAGAAATAATCGTGAGGCATTCCCACA |
| 26 | 21[98] | 16[105] | TTAGGAGCACTAGCATATCAA |
| 27 | 1[252] | 5[258] | TCAGAGCATAGGAAGTACAAATCATAGTCAGACGT |
| 28 | 20[69] | 21[77] | AGGATGCTCGAAAGGAATTGAG |
| 29 | 15[147] | 3[146] | TACATTTATAATTATTTGCACTATCAAACCCTTAGTAATGGT |
| 30 | 9[70] | 23[69] | CGCCAGATTACATTATTGCAACCCAGTCACGACGTATTTAGA |
| 31 | 5[315] | 14[322] | AAAGGTGGCATCAAAAATCATAATAAAGGAGAGTC |
| 32 | 10[111] | 22[112] | ACTATCGATTTACAAAATCGCGATGTGC |
| 33 | 20[300] | 15[314] | GGAACAAGATTTAGGAATAGCGTCTGGCCGGATTCGCTTTTG |
| 34 | 6[300] | 20[301] | GATTGCACAAATATCGCGTATAGAGCAAGGCTTTTTCCTAAC |
| 35 | 17[203] | 0[196] | GCAAGGCAGAATCAATGATACCAGTAAGAGCGGGGAGCCCGGAATAGGT |
| 36 | 10[300] | 18[294] | TTCATTGCAATACTTTTGCCAAGCGAGACACTATCGAATTACAAAACTA |
| 37 | 10[279] | 22[280] | TCGTCACATGTTTAGACTGGAGGACGAC |
| 38 | 16[104] | 19[111] | ACCCTCACACCTTGTGGCCAAGTTAGAATCAGATACCACACCGCGGTCA |
| 39 | 19[91] | 21[97] | AAAACAGACAGTGCAAAGCATATCAATAAATATCT |
| 40 | 0[230] | 10[217] | CGAGAGGGTTGATACACCCTCGAGGCAGCCGCCGCTCAGAGC |
| 41 | 21[266] | 16[273] | AGTAGTAAATTGAAGGATATT |
| 42 | 23[140] | 12[147] | GTTTGAGTCGCTATCAATTACCTGAGATTACGGGA |
| 43 | 11[168] | 19[174] | TTTAGCGATCAGATTTTATTTGGTATTAAAACCAACTGAACAGAGTTAA |
| 44 | 6[69] | 20[70] | AGGAATGTTTGCTTTGACGAGCTGAAAGGATTCACACATTTG |
| 45 | 0[314] | 8[301] | ATTAGGCTGAATATAGTTTCATTAGAAT |
| 46 | 11[84] | 19[90] | ATAACATCCATCACGAAGTGTCCGATTAAACGTGCCGCCGCTAGAAGAT |
| 47 | 19[175] | 21[188] | GCCCAATCTATCTTAGCCGAAATAATAATAGCAAACGTAGAA |
| 48 | 21[224] | 14[231] | CATTTGGGGAGGGATCGGTCATAGCCCTACATAGC |
| 49 | 9[273] | 0[266] | TACCGTTAGTCTCCAAAAAAAGCCTGTAGGATAGC |
| 50 | 23[308] | 12[315] | TGGGAACTCGTAACCGTCAATATGATATGAGGGTA |

| | | | |
|---|---|---|---|
| 51 | 11[126] | 19[132] | AACAATTACATAAATAAATCGGAAAACAGTCAATATACCTTTGGTAATT |
| 52 | 4[48] | 21[55] | AACAGCTCCCTAAAACATCGCTTTGAATCTGTAAGTCGCCCTAACCCCG |
| 53 | 22[111] | 9[104] | TGCAAGGTTAAATCGAACAACTAATAGAAAGGGACTGACGCTATCAGTG |
| 54 | 12[195] | 23[202] | CGGAACCTTGCGTCAGACTGTATTCATTCAGCGCC |
| 55 | 14[111] | 18[105] | ATTTTTCCTGAACCTAACACC |
| 56 | 0[265] | 10[259] | AAGCCCACACCACCACAATGAGGAGTTAAGCGAAA |
| 57 | 22[132] | 16[126] | GCTGGCGTATTAATGAACAAAATTCCTGCCTGATT |
| 58 | 12[314] | 0[315] | GCTATAATAATACTCATATATTTCCATATAACAGTTTTGACC |
| 59 | 17[119] | 0[112] | GAATAATAAGAAATGAGAAGATAGCGATTTTCATCTTAAATAAGAATAA |
| 60 | 18[188] | 2[182] | GCAATAGAATAAGAACAATAGTAATGCAAATATAA |
| 61 | 9[154] | 23[153] | ACAAGCAACAGCCATTTATCCCGGTGCGGGCCTCTTAGAATA |
| 62 | 1[280] | 10[301] | AACTAAAAATCAGGGGATGCTTTAAACAGTTCAGAAAACGTAATAAATA |
| 63 | 0[111] | 12[112] | ACACCTGGGTATTTTAGTTAAAGCTTAGTAACCTTTTGAATA |
| 64 | 8[300] | 22[301] | GACCATATAGTCAGAAGCATGAAGAAGTGCGGAATCGTGCAT |
| 65 | 20[187] | 6[189] | AAAGGTTTTTTGTAGAGCCTAGGAATCATTACCACACCTTA |
| 66 | 10[258] | 18[252] | GACAGCATTGAGGAAACGGGTCTAAAACTCTTTGAGACCTTCTACAGAC |
| 67 | 11[315] | 19[321] | TGCCTGAGAACCCTTTTGCGGAAAAACAATTAAGCACAGGCAAAAACAG |
| 68 | 15[315] | 3[314] | AGAGATCTGGAGCAAACAATATAGCAAATTATGACTTATACA |
| 69 | 17[147] | 17[146] | ATCAAAACATAAAGGAACTGGCATGAAATTCATCACTACCAT |
| 70 | 5[259] | 23[265] | TAGTAAATTTCAACAATTTTTACACTCAGAAAGAGTTTAATTATACCAG |
| 71 | 22[174] | 16[168] | TTGGGAACAATCAATATAAAAAGTATGTCGGAATA |
| 72 | 2[223] | 22[217] | GAAAGTACAGTACCTGAATTTAAGCCAGTTTGCCACTCAGAGGTGCCGG |
| 73 | 21[119] | 1[125] | TCATCATGAAACCACAGATGAGATTGCTGCTTCTGTCAATATACTAGAA |
| 74 | 0[97] | 10[91] | GGTGCCGCCCAAATAGTCTGTCACTTGCTGGTAAT |
| 75 | 10[216] | 18[210] | CACCACCCGCCTCCTCTTTTCTTTCATCTAGCGACCGGAAACTCATCGC |
| 76 | 2[139] | 22[133] | AAGACAACTAAATTAATCCTTTCGCTATTAACGGATTCATTTTACGCCA |
| 77 | 14[195] | 18[189] | CTTTAAAACCAGAATGAAATA |
| 78 | 16[272] | 19[279] | CATTACCGTAATCTCACTAAATCACGTTGAAAAGATTTTGCTGACCAAC |
| 79 | 0[160] | 9[153] | AACGCCAACAAATTGCTTATCCGGTATTTTCGAGA |
| 80 | 2[55] | 22[49] | AGCAGGCAAAGAATCATTAATGTCGGGACTACGTGCGTAATCCGACGGC |
| 81 | 2[244] | 7[237] | ACGATCTGAGTTTCATTGTATCGGTTCGTTGAACA |
| 82 | 21[203] | 1[209] | ATCACCGGGAAATTAGCGCGTATAATCAATAAATCGATTGGCAGGTTTA |
| 83 | 23[182] | 11[188] | TTCATATAGGCTGCAGCGTCTTAAATCACGACTTG |
| 84 | 23[98] | 8[105] | ACTCGTACGATTAATACCTACGCCTTGCCTGAGTAGAAGAACTGACCGA |
| 85 | 22[216] | 16[210] | AAACCAGAAAGGGCTATTGACTCACCGATCACCAG |
| 86 | 9[322] | 22[322] | GAGAAGCTGCCGGATCAACCGTGTAGAT |
| 87 | 11[210] | 19[216] | ACCAGAGGTCAGACCTCATTAACCGTTCAGGAGTGTTGAGTAGACCTGC |
| 88 | 1[112] | 11[125] | CATAATTATGTGAGTCTTAATTACATTT |
| 89 | 9[238] | 23[237] | GCTTGCTGTAATGCCGGCTACAGCTTTCCGGCACCTGTATGC |
| 90 | 11[70] | 11[69] | TTCTTTGTACCGCCAGGTTTCCTGTGTGTGTAAAGCCAATAC |
| 91 | 22[300] | 23[307] | CTGCCAGTAATAAAACGAACG |
| 92 | 0[244] | 9[237] | ACCGTAACAGAACCCACGCATAACCGCAAGTATCA |
| 93 | 21[161] | 1[167] | ATTACGCGAAACGCCCAATCCAAAATAAAGCCGTTATAGAAGCTTACCA |
| 94 | 7[189] | 19[202] | TCTTTTGTTTATCAGCAAGAAACGTCGA |
| 95 | 6[188] | 0[182] | TCATTCCAGTAATAGGCTTAA |
| 96 | 23[154] | 11[167] | AGTTTATTTTGTCAGGGCGATTGAATCTTTTGCACGAGGCGT |
| 97 | 23[70] | 11[83] | AGTATTAGACTTTAGGGTTTTCAGGAAAACAATATATTAGTA |
| 98 | 0[76] | 9[69] | GGAACCCGGCCCACAACCGTTGTAGCTGCGCGGTA |
| 99 | 5[133] | 23[139] | TATATGTGAGAGACGTGAATTGTAAAACGTAACAGTTTTGCGTTTAAAA |
| 100 | 1[84] | 5[90] | ACCATCATAAAGCATAGAGCTGGAAGGGGGCAAGT |

| | | | |
|---|---|---|---|
| 101 | 23[203] | 10[196] | AAAGACAGCAAAGCCCGGAACCTCAGAGCCGCCTT |
| 102 | 22[90] | 16[84] | AACGCCACAAACAACGTCAATTATCTAATCTGGTC |
| 103 | 12[62] | 0[63] | GCTTAAGAACCTGTGGGTGCCTCTATCAGGGCGATTAGGTTG |
| 104 | 5[203] | 5[202] | AGTTAATGCTGAGACTCCTCAAGAGAAGGAACGCGCCTAAAC |
| 105 | 19[112] | 14[112] | GTACAAAGTCAGAGTTAACCTCCGGCCAAGACGCTTGCGTAG |
| 106 | 21[56] | 14[63] | CTTCTAAGCACGACTGTAATGAGTAACACGGGGCA |
| 107 | 2[160] | 6[154] | AATTCTGGTAATTTTACCGCA |
| 108 | 18[125] | 17[118] | TGAACACCCTGAATTCCTTCT |
| 109 | 3[315] | 6[301] | TTTCGCATTTGGGGCGCGAGGATTAGAGAGCTTAACCAAGCG |
| 110 | 4[286] | 18[273] | CGAACGAAGCCCGATCGTTTACCAGATATGACAAGAAAGAGG |
| 111 | 23[266] | 11[272] | TCAGGACTCGGCCTCTTTTTCCCTCAGCAAGGCCG |
| 112 | 11[189] | 1[188] | CGGGAGGTTCGCGCCCAATAGCTCAACA |
| 113 | 5[49] | 23[55] | TTGCAGCGACGGGCGGGTGGTAGGGGCCTGGTTGGTTAAGTGGTGGTTG |
| 114 | 1[168] | 5[174] | GTATAAATCGCCATGGCATTTACAAAAGACAACAT |
| 115 | 16[237] | 16[238] | AGTGAATTAGAGCCATGAAACCATCGCAAGCGCAGCTCATTC |
| 116 | 5[175] | 23[181] | GTTCAGCATAAGTCTCAATAAAATGAAAAAACGATGGCAACATAGAAAA |
| 117 | 18[209] | 17[202] | CTGATAAATTGTAAGGCATTA |
| 118 | 5[217] | 23[223] | GCCTATTCAGTGCCTACTGGTTAATCAGGGCATTTAGGTAAAGACATTC |
| 119 | 23[224] | 12[231] | AACCGATGCTTCTGCCGCCACCCTCAAACACCCTT |
| 120 | 12[111] | 23[118] | CCAAGTTATCAGGTTTAACGTCCAGAAGCTTTGCC |
| 121 | 17[315] | 5[314] | TCATTCGTTAAAGATGAACGGAGCCCCAAGGCAAACAGCTGA |
| 122 | 9[105] | 0[98] | AGGCCAAATGCGGGAGCTAAAGAAAGCCGGGTCGA |
| 123 | 16[69] | 16[70] | CAGTTTATTTACGCCAACTCGTCGGTTAAGAATAGAAATCAA |
| 124 | 23[287] | 10[280] | TCTACGTTTTGAGGTAGCGTCAATCCCCCTCAAGA |
| 125 | 14[230] | 2[224] | AGCACCGAATAAGTCAGTCTCAGGCGGATAAGTCTAAATTCTGAAACAT |
| 126 | 18[293] | 17[286] | GCATGTCAATCATGAATAACG |
| 127 | 18[104] | 2[98] | GCCTGCAAGGTGAGCGCCGCGTCACGCTAACGTGG |
| 128 | 21[245] | 1[251] | CTGACGATAATCATCACCAACAAAATACTTCGAGGCATCGCCGCCACCC |
| 129 | 1[189] | 11[209] | GTACCGTACTCAGGCTTGATATTACCAGAACCACC |
| 130 | 16[251] | 0[245] | AAAGCTGGCTGGCTCCCCCAGGAATTGCGCCTTTAGTCACCACCCATGT |
| 131 | 10[237] | 10[238] | GCGAACCGCCACCCCAGCATTGAATATATTCGGTCGAGGGTA |
| 132 | 10[132] | 18[126] | AAACATCCGAATTATTCGCCTATATACAAGAAATAGGAAGGGAATTAAC |
| 133 | 16[300] | 21[286] | TTCAACTAATGCAGATACACCGATTCAT |
| 134 | 18[251] | 2[245] | CAGGCGCAGACGGTGCGGAGTGTCTTTCTAGCGTA |
| 135 | 10[174] | 18[168] | TTGCTATTACCAACCAGTTACAAATAAGATAGCAGAGCAGATACCGAAG |
| 136 | 12[146] | 0[147] | GAAACAATAATTAATTAATGGATCATATGCGTTATACATACC |
| 137 | 10[90] | 18[84] | ATCCAGAAACGCTCCTGAAATACGACCAAGAACCCAAAATCTCACGCTG |
| 138 | 19[203] | 14[196] | AATCCGCACAGTGCCCGTATGGCTTTTGAGTTTGC |
| 139 | 18[237] | 5[237] | AAAGTACAACGGAGTACTTAGACGGGGTTCGGAACCTATTAG |
| 140 | 8[104] | 1[111] | GTAAAAGCAAGTTTTTGGAAT |
| 141 | 11[154] | 11[153] | AGAACGCCCAGCTACACAAAAGAAGATGAATTACCTTTTCTA |
| 142 | 16[167] | 0[161] | CCCAAAAAAAAGTACCTTTACCATGTAGAACCAAGAGGCAGAATTTAAC |
| 143 | 19[280] | 4[287] | TTTATGTACCCCGGTTCAAAG |
| 144 | 11[273] | 1[279] | CTTTTGCTCATAGTTGCGCCGCTCATTTTCTATGC |
| 145 | 12[279] | 23[286] | AAAATGAAACGACGATAAAAAACAGGTAAGAAAAA |
| 146 | 0[279] | 12[280] | TTAAAAGGCAGAGGTCATTTTGATTAAGTACCCTGTAATAGT |
| 147 | 18[83] | 2[77] | AGAGCCACACCAGCACAGGGCGGGCGCTAAGAAAG |
| 148 | 14[62] | 2[56] | CGAATATTTTTCTTCCAGCTGAGCCCGAGATAGAAAGCTGGTTTGCCCC |
| 149 | 10[321] | 11[314] | GTCAAATCACCATCAATGCAA |
| 150 | 11[252] | 19[258] | CTTGCAGCAACAACTGAATTTAAAAGGAGAATAATAGTTTCACAATCAT |

| | | | |
|---|---|---|---|
| 151 | 1[294] | 4[301] | GTCTGGAAATGCTGGGCTTAGAGTACCTAAACTCCAACAGGTGATTTAA |
| 152 | 21[287] | 1[293] | CAGTTGACATTATTCCAAAATGAGGGGGACTATTAAATCAAAGTACGGT |
| 153 | 23[56] | 12[63] | TGAATTCTGTAAAAATGGTCATAGCTCCGGACAGG |
| 154 | 23[238] | 11[251] | GATTTTAAGAACTGTCCAGCCAGAGGCTTCGGAACGCTGAGG |
| 155 | 12[230] | 0[231] | ATTAGCGAATGGAAGGAGGTTAGAACCGCCACCCTCAGCCGT |
| 156 | 16[209] | 21[202] | TAGCACCATTACAAGTGAATT |
| 157 | 18[272] | 2[266] | ACAGATGCCGAACTAAACAACTGAATTTTCCACAG |
| 158 | 5[147] | 4[154] | GATGCCAGACGACGTAATATC |
| 159 | 2[97] | 22[91] | CGAGAAATGACGGGCAGGAGGTTTTATACAATCGTATGGAAAGTTGGGT |
| 160 | 16[125] | 21[118] | GTTTGGATTATAAAGGAATTA |
| 161 | 5[91] | 23[97] | GTAGCGGCTTAATGTTTCCTCCAGAGATGTAATAATTAGAGCTTCGACA |
| 162 | 23[119] | 10[112] | CGAACGTAAAGGGGGCAGAGGAAGAAAACAAAAAA |
| 163 | 4[153] | 18[154] | CCATCCAGAGAGATAACCCACTAAAACA |
| 164 | 0[146] | 10[133] | GACCGTGTGATAAAGTTTAGTAAACAGTTCATTTGATGAAAC |
| 165 | 22[321] | 17[314] | GGGCGCAAAACGGCAACCCGTCTTCCTGCCATCAAAAATAATAGGAGAA |
| 166 | 19[133] | 21[139] | GAGCGCTACGGGAGTTAGAACATATAATATTATCA |
| 167 | 2[76] | 6[70] | CGAAAGGGGGAGCCTTTAGAC |
| 168 | 18[69] | 5[69] | GCACAGACAATATTCATTAAACCAGTGAAAGCGGTCCACGCG |
| 169 | 8[41] | 5[48] | ACTCACAACGTCAAAGGGCGATTGGAACTAAATCAGAAAATCGAGAGAG |
| 170 | 19[259] | 21[265] | AAGGGAAAACGGTGATCAAGACAAATCACAGAACG |
| 171 | 5[119] | 5[118] | TTGGGTTGAGAAAACTTTTTCAAATACGGCGCGTAACTTAGG |
| 172 | 0[62] | 10[49] | AGTGTTGTTCCAGTAAAACCGTAATGAGCATAAAGAAATTGT |
| 173 | 18[153] | 5[146] | GGGAAGCGCATTAGAATATCATAGGTCTAAATGCT |
| 174 | 8[348] | 1[345] | ttttttttGCAAGGATAAAACAATTCTGCtttttttt |
| 175 | 5[22] | 2[15] | ttttttttCCGCCTGGCCCTCTGTTTGATGGTGGTTCCGtttttttt |
| 176 | 9[18] | 12[15] | ttttttttTTGCGCTCAGATAAAGACGGAtttttttt |
| 177 | 14[345] | 7[348] | ttttttttTCATTGCCTCCTCAGAGCATAtttttttt |
| 178 | 1[15] | 8[18] | ttttttttACGTGGACTCCATTAATTGCGtttttttt |
| 179 | 6[41] | 0[15] | CCAACGCTCCCTTAAAGAGTCCACTATTAAAGAtttttttt |
| 180 | 20[348] | 15[345] | ttttttttTTAAATGTGAGCGCTATCAGGtttttttt |
| 181 | 3[15] | 6[18] | ttttttttAAATCGGCAAAAGCGGGGAGAtttttttt |
| 182 | 6[348] | 3[345] | ttttttttAAGCTAAATCGGAATAACCTGtttttttt |
| 183 | 12[345] | 9[348] | ttttttttATAAATTAACTTTATTTCAACtttttttt |
| 184 | 13[15] | 22[18] | ttttttttGGATCCCCGGGTCTCAGGAGAtttttttt |
| 185 | 21[18] | 9[41] | ttttttttTTATGACAATGTCCCGTGCTGAAGAACTCTTTACCTCCTGCCCG |
| 186 | 15[322] | 23[348] | TACAAAGGAGTAACGGATTGACCGTAATGGGATtttttttt |
| 187 | 23[18] | 10[22] | ttttttttAGCCAGGGTGGATGTTAAGCTTTACCGAGCTCACAATTCCACtttttttt |
| 188 | 11[22] | 7[48] | ttttttttACAACATACGAGCCGGAAGTGAGCTACTTTCCAGAATCGGGGCGCCA |
| 189 | 0[345] | 10[322] | ttttttttGAACGAGTAGATTTAGTGATTCCATTTTTAGTAATGTGGAGACA |
| 190 | 22[348] | 13[345] | ttttttttAGGTCACGTTGGTTCTAGCTGtttttttt |
| 191 | 22[48] | 16[22] | CAGTGCCCTTCTAATCCTTAGCCAAAATGGAGTGACTCTATGATACCtttttttt |
| 192 | 19[22] | 4[22] | ttttttttGCGAACTGATAGGATTGCCCTTCAtttttttt |
| 193 | 17[42] | 18[22] | CTGCCATGGCTATTAGTCTTTAATGCtttttttt |
| 194 | 7[18] | 14[15] | ttttttttGGCGGTTTGGCATTTCACATAtttttttt |
| 195 | 19[322] | 21[348] | GAAGATTTATTTTGCATTAAAAGGAACGTAGCCAGCTTTCATCAACAtttttttt |
| 196 | 15[15] | 20[18] | ttttttttAATCATTTCTCCTTGTCAACCtttttttt |
| 197 | 17[22] | 7[41] | ttttttttGACAGTGCGGCCGGCTGACCGTATTG |
| 198 | 2[345] | 6[322] | ttttttttTTTAGCTATATTTTCAAATGGTCTTGTACC |
| 199 | 18[352] | 19[352] | ttttttttAAATTGTAAACGTTAAGTATAAGCAAATATTTtttttttt |
| 200 | 10[352] | 11[352] | ttttttttAAGGGTGAGAAAGGCCGTAGGTAAAGATTCAAtttttttt |
| 201 | 16[352] | 17[352] | ttttttttTCATTTTTTAACCAATTTTTTGTTAAATCAGCtttttttt |
| 202 | 4[352] | 5[352] | ttttttttAGCATTAACATCCAATTTCTACTAATAGTAGTtttttttt |
